# An enhanced yeast display platform demonstrates the binding plasticity under various selection pressures

**DOI:** 10.1101/2020.12.16.423176

**Authors:** Jiří Zahradník, Debabrata Dey, Shir Marciano, Gideon Schreiber

## Abstract

Yeast surface display is popular *in vitro* evolution method. Here, we enhanced the method by multiple rounds of DNA and protein engineering, resulting in increased protein stabilities, surface expression, and enhanced fluorescence. The pCTcon2 yeast display vector was rebuild, introducing surface exposure tailored reporters – eUnaG2 and DnbALFA, creating a new platform of C and N terminal fusion vectors. In addition to gains in simplicity, speed, and cost, new applications were included to monitor protein surface exposure and protein retention in the secretion pathway. The enhanced methodologies were applied to investigate *de-novo* evolution of protein-protein interaction sites. Selecting binding from a mix of 6 protein-libraries towards two targets using high stringency selection led to the isolations of single high-affinity binders to each of the targets, without the need for high complexity libraries. Conversely, low-stringency selection resulted in the creation of many solutions for weak binding, demonstrating the plasticity of weak *de-novo* interactions.

## Introduction

Macromolecular interactions are a driving force of most life processes. Proteins bind fast and specific even in the crowded environment of the cell, transferring signals, building complexes, transport cargo, and much more. This happens in an incredible range of concentrations, from millimolar to femtomolar. The generation of novel, specific interactions has been a major goal of protein engineers from the beginning. For example, the generation of novel antibodies binding specific targets has revolutionized medicine, as acknowledged by the 2018 Nobel prize in chemistry, which was awarded for the development of the phage display method for *in vitro* evolution of antibodies to specifically bind any given target. Phage display was the first of many other *in vitro* evolution methods since devised. In 1993, the pioneering work of Schreuder and colleagues (Schreuder et al., 1993), first described yeast display, which over time became the most widely used method for directed protein evolution. Similarly to other display methods, its principle is based on cycles of naïve protein library exposure, selection, and enrichment of yeast clones with desired properties. Yeast display has proven to be an effective method for developing, improving, and altering activities of proteins for research, therapeutic, and biotechnology applications. The unprecedented power of the technique, together with its relative ease of use and reasonable cost has made it popular in many laboratories around the world. The use of *Saccharomyces cerevisiae* and its homologous recombination machinery reduces the need for laborious DNA library preparations, with only DNA fragments being needed (Benatuil et al., 2010; Swers et al., 2004). Coupling of the genotype-phenotype association with high-throughput single-cell analysis on a fluorescent activated cell sorter (FACS) offers a simple and efficient screening process, with low risk of false-positive results (Chao et al., 2006).

The most popular yeast display setup is based on the yeast mating factor agglutinin A protein, which is composed of two independent domains: the A-agglutinin GPI-anchored subunit (Aga1p) and the adhesion subunit (Aga2p) (Schreuder et al., 1993). The subunit interaction is mediated by two disulfide bridges and the protein of interest is fused to the plasmid-encoded Aga2p subunit (Boder and Wittrup, 1997). Both C and N terminal fusions with Aga2p were used for successful display on the yeast surface (Uchański et al., 2019; Wang et al., 2005). Multiple Aga2p fusion partners, and their libraries, were screened and tailored to fulfill a plethora of tasks such as affinity reagents development (Gai and Wittrup, 2007; Simeon and Chen, 2018a; Mata-Fink et al., 2013), substrate specificity modulation (Cohen-Khait and Schreiber, 2016; Zhang et al., 2013), protein stability engineering (Traxlmayr and Obinger, 2012), and also for directed evolution of enzymes (Mei et al., 2017; Szczupak and Alfonta, 2015). Still, despite all of these developments, yeast display selection of optimal binders is challenging and time-consuming.

In the first part of this manuscript, we aimed to optimize all steps of the methodology for display and selection. In the second part, we tested the new method for a challenging biological question: how easy is it to evolve binding sites between random proteins that naturally have no connection between them, and probably never saw each other. This is a fundamental question in biology, as specific interactions have to compete against a vast majority of potential, non-relevant weak interactions with non-cognate partners. These may suppress orderly biological functions within the proteome (Schreiber and Keating, 2011). We have previously used conventional yeast display to approach this question for the interaction between the TEM1 β-lactamase and its protein inhibitor BLIP, showing that many interface solutions maintain this complex (Cohen-Khait and Schreiber, 2016). Moreover, we have shown that a TEM1 heterodimer (TEM1_WT-TEM1_mutant) could be evolved by incorporating only two mutations (Cohen-Khait 2017). In this manuscript, taking advantage of the optimized yeast display method, we were able to obtain a much broader picture on the evolution of high and low-affinity binders between unrelated proteins, with major implications to biology.

### Design

We aim to evaluate the emergence of new protein-protein interactions among 6 different protein libraries and 3 different prey proteins with low and high stringency selection. From this, we will learn about the easiness of creating new protein-protein interactions between random pairs of proteins. Moreover, using low and high stringency selection, we show what distinguishes low from high-affinity interactions in evolutionary terms. To make this possible we looked for substantial simplification of the current yeast surface display methodology. The graphical abstract provides a summary workflow for this project. In the first step, we used restriction-free cloning for complex plasmid backbone minimization of the original pCTcon2 vector, to remove all non-necessary parts and thus making it much shorter (Figure 1, step 1). Next, we screened for different reporter proteins and evaluated the efficiency of their yeast surface exposure, which is essential for display. The best reporters were subjected to multiple rounds of protein engineering to optimize their yeast surface expression and brightness (Figure 1, step 2). Then, we optimized signal peptide and protein linkers to allow for both N and C terminal library insertions (Figure 1, step 3). The performances of the newly developed plasmids were assessed by quantifying the yeast surface expression levels of 16 different proteins (Figure 1, step 4). After system optimization, enhanced yeast surface display was evaluated for parallel high and low-stringency selection for 6 different scaffold protein libraries. The aims here were to evaluate the relation between selectability and protein stability (Figure 1, right panel), and to assess the number of parallel binding solutions with respect to binding affinity.

**Figure 1.**
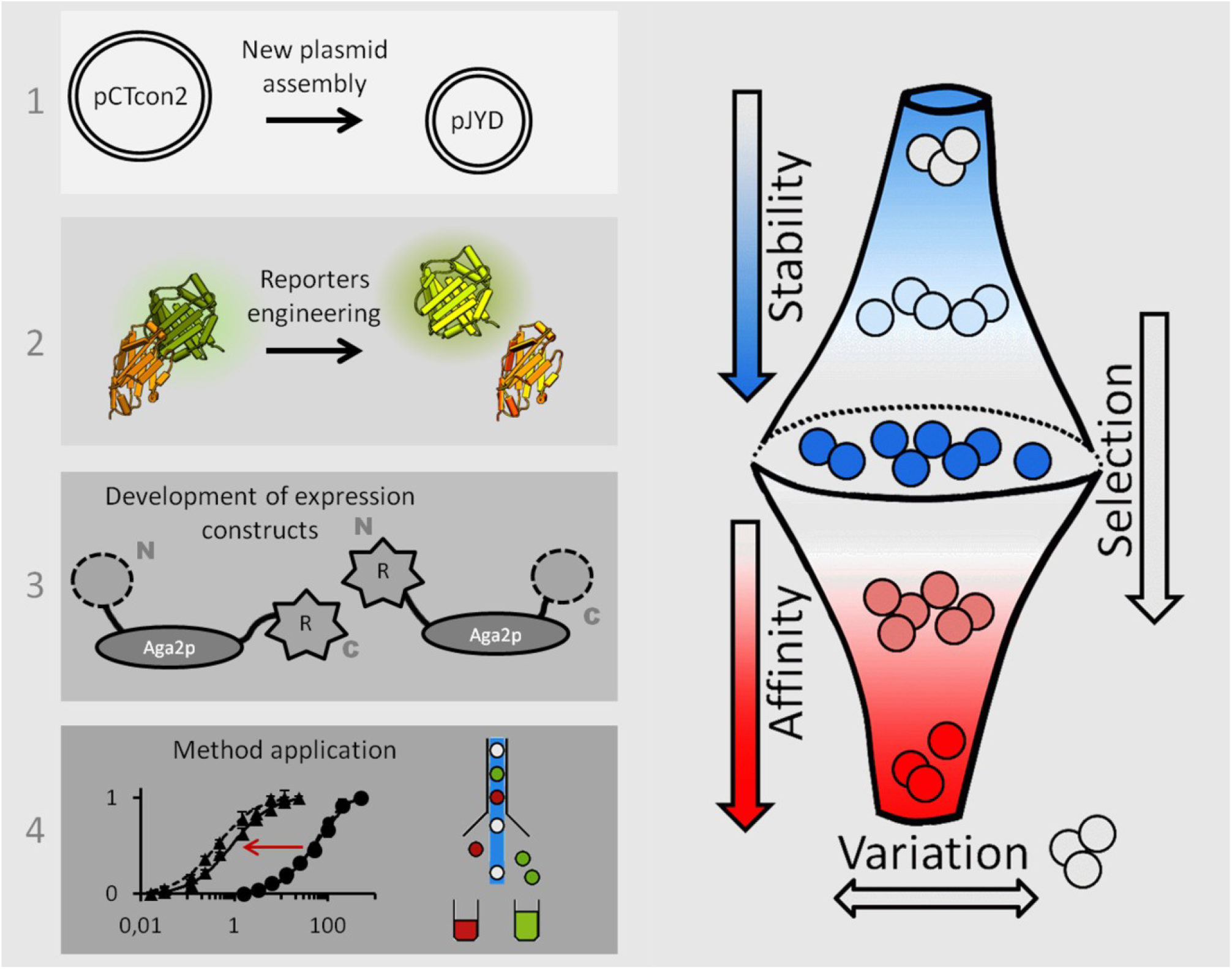
Experimental workflow and expected outcomes.

## Results

### Part I: optimization of the yeast display platform

#### Yeast display plasmid optimization

The most frequently used plasmid for yeast display, pCTcon2, was developed more than 15 years ago (Colby et al., 2004), before advanced DNA manipulation technologies such as restriction-free cloning (van den Ent and Löwe, 2006) were developed. To simplify the work with pCTcon2, we modified its backbone to create a new plasmid system. First, we replaced the *AmpR* gene with *KanR* coding for aminoglycoside-3’-phosphotransferase, as kanamycin resistance is more stable over time. Next, we removed multiple unnecessary sequences, like T7, T3 promoter regions, F1 origin of replication, lac operator, and promoter fragments, which unnecessarily increased its size. We used three components assembly by restriction-free cloning (Peleg and Unger, 2014) to reassemble a new plasmid lacking these sequences. The mutual position of functional elements was kept the same as in pCTcon2. The resulting vector was designated pJYD (plasmid J-series yeast display) and its full-length sequence was verified. pJYD is 1301 bp shorter than the parental vector pCTcon2 (6456 bp).

#### Screening for expression reporters

A key component for antibody-labeling free yeast display is an efficient reporter protein system with low risk of false-positive readings. We tested a broad range of four green, five far-red fluorescent proteins, and two nanobodies of different sizes and properties. The two colors were chosen to fit the most popular FACS fluorescence setup, corresponding to green and red channels FL1 and FL4 respectively, while allowing tests such as propidium iodide viability staining (Davey and Hexley, 2011). We examined the FACS expression characteristics of green fluorescent proteins mNeonGreen (Shaner et al., 2013), yeast enhanced green fluorescent protein (yeGFP) (Huang and Shusta, 2005a), bilirubin-inducible fluorescent protein UnaG (Kumagai et al., 2013), FMN-inducible fluorescent protein iLOV (Chapman et al., 2008), biliverdin-binding far-red fluorescent proteins dFP-mini (Sheehan et al., 2018), GAF-FP (Rumyantsev et al., 2015), TDsmURFP (Rodriguez et al., 2016) with two Y56R mutations (Fuenzalida-Werner et al., 2018) IFP1.4 (Shcherbakova and Verkhusha, 2013) and miRFP670nano (Oliinyk et al., 2019). In addition, we tested two peptide tags recognizing nanobodies for their performance and possible uses in yeast display to extent labeling strategies. We used BC2 peptide-tag nanobody (Braun et al., 2016) and ALFA-tag binding nanobody (Götzke et al., 2019). The codon-optimized genes (Kaishima et al., 2016) were cloned into the pJYD plasmid in two different positions to obtain plasmids with protein expression at different localizations - cytoplasmic expression and cell surface expression, fused with Aga2p. Both positions were under the control of the GAL promoter. The expression media was supplemented either with 2.5 nM bilirubin, 1 mM FMN, or 2.5 nM biliverdin for UnaG, iLOV, and far-red fluorescent proteins respectively. We compared fluorescent intensity differences after 16 hours of expression at 20 °C by flow cytometry. Results were further validated using fluorescence microscopy (Figure 2). The presence of characteristic inner ring for endoplasmic reticulum and Golgi or fluorescence foci suggesting the presence of fluorescent proteins in vacuoles were analyzed to uncover impaired reporter processing to the cell membrane (Drew et al., 2008).

**Figure 2.**
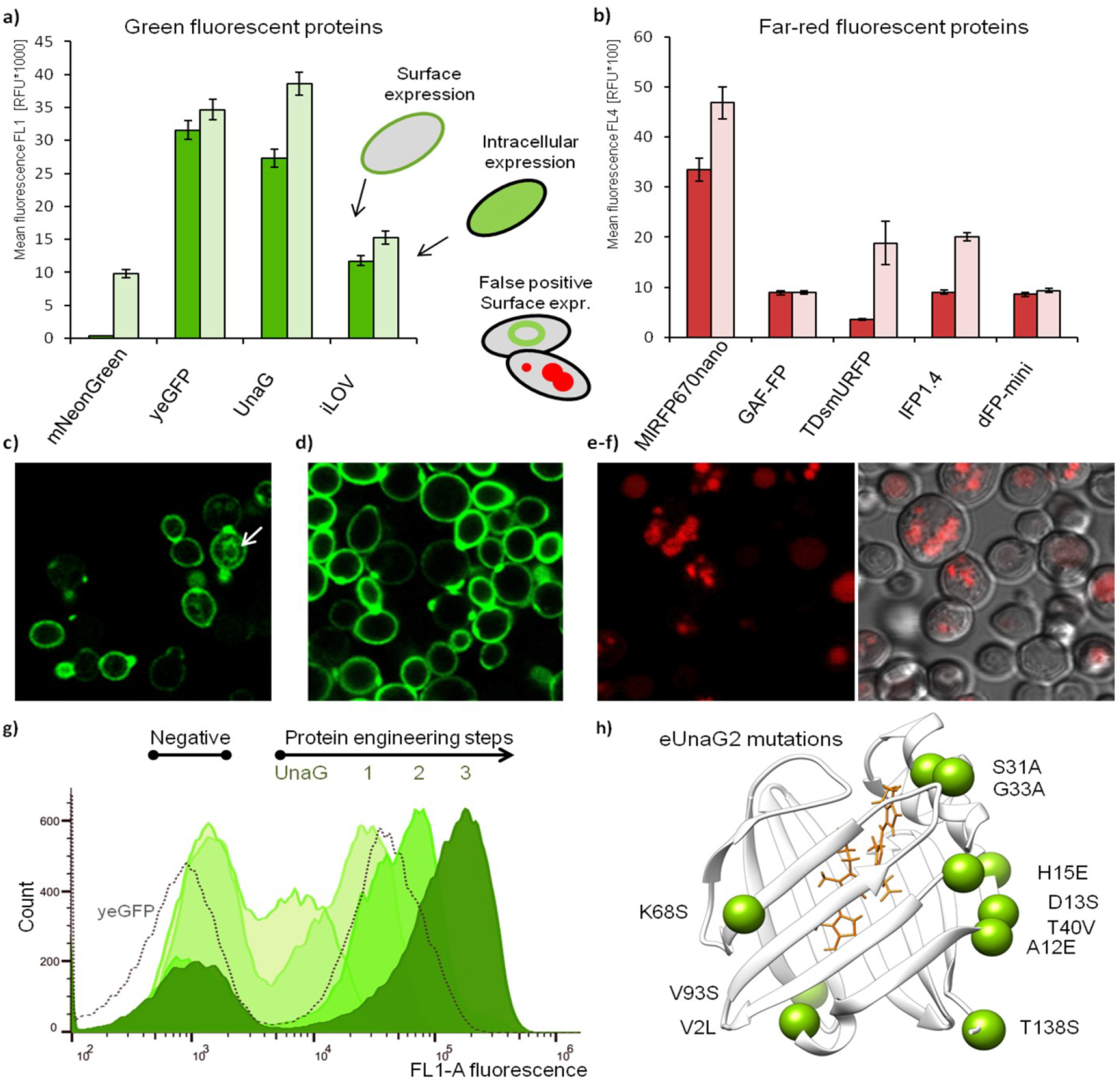
Evaluation and engineering of fluorescence proteins for optimal yeast surface exposure. a) Comparison of cytometry assessed mean fluorescence intensities for a) green and b) far-red fluorescent proteins between Aga2p fusion on the cell surface (rich color) and intracellular expression (faint color). c-f) Microscopy images of *Saccharomyces cerevisiae* EBY100 cells expressing c) yeast enhanced yeGFP; False-positive signal from yeast endoplasmic reticulum is marked by the white arrow. d) UnaG bilirubin dependent fluorescent protein and e-f) miRF670nano protein. In contrast to yeGFP and UnaG, the miRF670nano protein fused to the C-terminus of Aga2p was not detected on the yeast surface. g) Flow cytometry histograms showing the green fluorescence signal (FL1 channel) distribution among cell populations during the subsequent protein engineering steps of UnaG. The dotted line shows yeGFP protein for intensity and distribution comparison. h) Mutations introduced during the eUnaG2 protein engineering depicted in the 3D structure of UnaG protein (PDB id: 4i3b).

Our cytometry results showed the highest fluorescence intensities of surface-exposed protein for the yeGFP construct, while UnaG showed higher cytoplasmic fluorescence levels. Validation by microscopy of yeGFP showed a higher proportion of fluorescent signals emitted from the endomembrane than observed for UnaG (Figures 2c, 2d). Folding of fluorescent proteins inside the endoplasmic reticulum or other endomembrane compartments can be a source of false-positive signals in yeast surface display and should be avoided or decreased. Based on these results, its smaller size, and the ability to control the fluorescence by the presence of bilirubin, we used the UnaG protein for further tailoring its properties to best fit our yeast display setup.

Among far-red fluorescent proteins, only miRF670nano showed a satisfactory fluorescence signal on flow cytometry (Figure 2b). The microscope showed most of the recorded signal coming from inside the cells (Figure 2e, 2f). By the appearance of miRF670nano inside fluorescence foci, we hypothesize its localization to be mainly vacuolar. Therefore, we decided not to continue with biliverdin dependent far-red fluorescent proteins and instead focused on nanobodies. Both nanobodies showed high levels of expression on yeast surface when expressed on the C-terminus of the pJYD vector, with an ALFA-tag binding nanobody (nbALFA) being 24 % better in terms of total FACS signal. Based on these results, we decided to continue using nbALFA.

Next, we analyzed the ability of our reporter genes to be produced in an active – fluorescent form during the yeast cultivation in expression media. We tested whether the addition of bilirubin to the media will be sufficient for the UnaG fluorescent protein to be in its holo form without affecting yeast growth. At the range of concentrations tested (100 μM – 1 pM), we did not observe significant growth inhibition. Fluorescence saturation was observed with > 200 pM of bilirubin. Optimal labeling of yeast was achieved by using 1 nM bilirubin in the expression media added directly before the cultivation from frozen, DMSO diluted stock. This concentration led to a slight media color change. Since the stability of bilirubin might be compromised if unprotected from light at room temperature, the prolonged storage of the media with bilirubin is not recommended (Sofronescu et al., 2012). The expression labeling by using nbALFA during the cultivation was tested with different concentrations of ALFA-tagged mNeonGreen, produced in *E. coli* BL21. The stability of the fluorescent protein in media was routinely determined by measuring the fluorescence on a plate reader and did not decrease below 50 % of initial fluorescence.

#### Engineering UnaG for efficient cell surface expression and brighter green fluorescence

A high-quality fluorescence signal on the cell surface that reports expression requires bright fluorescence and a low level of the false-positive signal from inside the cell. Yeh et. al (Yeh et al., 2017) used yeast surface display to increase UnaG fluorescence, generating the eUnaG protein (enhanced UnaG). One single point mutation in eUnaG, V2L, led to an increase in thermal stability by almost 6 °C and doubling of the fluorescence intensity signal, suggesting that higher UnaG stability leads to increased fluorescence intensity. To further improve the protein’s stability, we combined computational and experimental procedures. Multiple stabilizing mutations were predicted by the consensual design of all available structures using the Pross web-server (Goldenzweig et al., 2016; Zahradnik et al., 2019). Since Pross was not designed to optimize secreted/surface-exposed proteins, we used the Pross suggested mutations as a starting point for random incorporation and selection, rather than testing suggested protein variants. A mutation library was created from the 13 *in silico* predicted mutations on EBY100 yeast cell surface. Cells associated with stronger fluorescence intensities were isolated by three-rounds of FACS sorting. Stronger fluorescence was verified for the isolated single clones. The brightest isolated clone eUnaG1 showed a significantly higher fluorescent signal when using FACS than the parental variant (Figure 2g, engineering step 1). eUnaG1 was further improved by incorporating additional 4 mutations found in other selected colonies (Figure 2g, engineering step 2). Finally, our inspection of the crystal structure PDB ID 4i3b showed a solvent-exposed hydrophobic patch formed between residues V89, V93, V98, V100, and V111. Since surface polarity is a critical parameter influencing expression (Magliery, 2015), we designed and tested the gain of N-glycosylation mutations (V93S, E107N) to rescind this patch. The gain of N-glycosylation was mediated by mutations leading to a new N-X-S/T surface-exposed motif (Gupta and Brunak, 2002). The V93S mutation led to a further doubling of fluorescence intensity (Figure 2g, engineering step 3). Altogether, we introduced 10 mutations in eUnaG2 (Figure 2h, which results in a 5-fold increase in its fluorescence intensity as measured by FACS. eUnaG2 expressed in *E. coli* BL21 cells, showed an increased thermal stability relative to UnaG of 10 °C (Figure S1b).

#### Engineering the ALFA-tag binding nanobody for efficient multicolor fluorescence labeling

The engineering of the ALFA-tag binding nanobody for increased thermal stability was a more challenging task, as computational tools for ΔΔG predictions of antibodies have higher false-positive rates (Baran et al., 2017; Goldenzweig et al., 2016). 10 mutations were predicted and tested one by one (Figure S1c). The proteins were expressed in *E.coli*, purified and their melting temperatures were measured using the nanoDSF Prometheus NT.48. We identified eight mutations with T_m_ values ranging from 54 °C (wild type) to 55.3 °C (best mutant Q69K) which is only 1.3 °C higher than the wild type. Combining all 10 mutations (see Figure S1c) lead to an increase of 6 °C in thermal stability. In a subsequent, second round of protein engineering, we screened two N-glycosylation gaining mutations (G17N, T25N) and their influence on the expression of the nanobody. Both of them slightly enhanced the protein expression, although decreased the thermal stability of the protein by 4 °C. The melting temperature of glycosylated proteins was measured directly on yeast by using the interaction with purified ALFA-tagged mNeonGreen (Traxlmayr and Obinger, 2012). Among all tested mutations, 10 mutations had positive effects on the recorded cytometry signals and were combined in the final construct of the protein, termed Designed ALFA-tag binding nanobody (DnbALFA). Figure S1d shows the development of the cytometry signal along with the protein-engineering steps. Measuring the binding affinity of DnbALFA and nbALFA towards ALFA-mNeonGreen showed a 2-fold reduction of the former (60 versus 25 pM respectively, Figure S1e). Despite the slight decrease in binding affinity, the gain in protein expression is much higher both on the yeast cell surface and in *E.coli* BL21 (DE3), showing a 10-fold increase in yield of the soluble designed protein (Figure S1f).

#### Yeast display platform design and engineering

In the next step, we incorporated eUnaG2 or DnbALFA either N or C-terminal to Aga2p, introduced the multi-cloning sites (MCS), tested signal peptides, and linkers to achieve optimal surface expression. Different plasmids were developed to create a complex platform allowing for a broad range of protein expression organizations and labeling strategies. An enhanced yeast display system has advantages in speed and variability of selection over the standard anti-myc antibody system. However, this variability requires quality control and careful experimental design. The following chapters are describing the plasmids’ development and their limitations. Further details are in the Supplementary material section, Step-by-step protocol.

#### Construction of N-terminal vectors

In the N-terminal fusion organization, the selected protein is bound N-terminal, between signal peptide and Aga2p, which is opposite to its location in the traditional pCTcon2 construct (Figure 3a) (Uchański et al., 2019; Wang et al., 2005). This construct has the advantage of the presence of a reporter gene at the C terminus avoids truncated protein expression and is thus less prone to artifacts.

**Figure 3.**
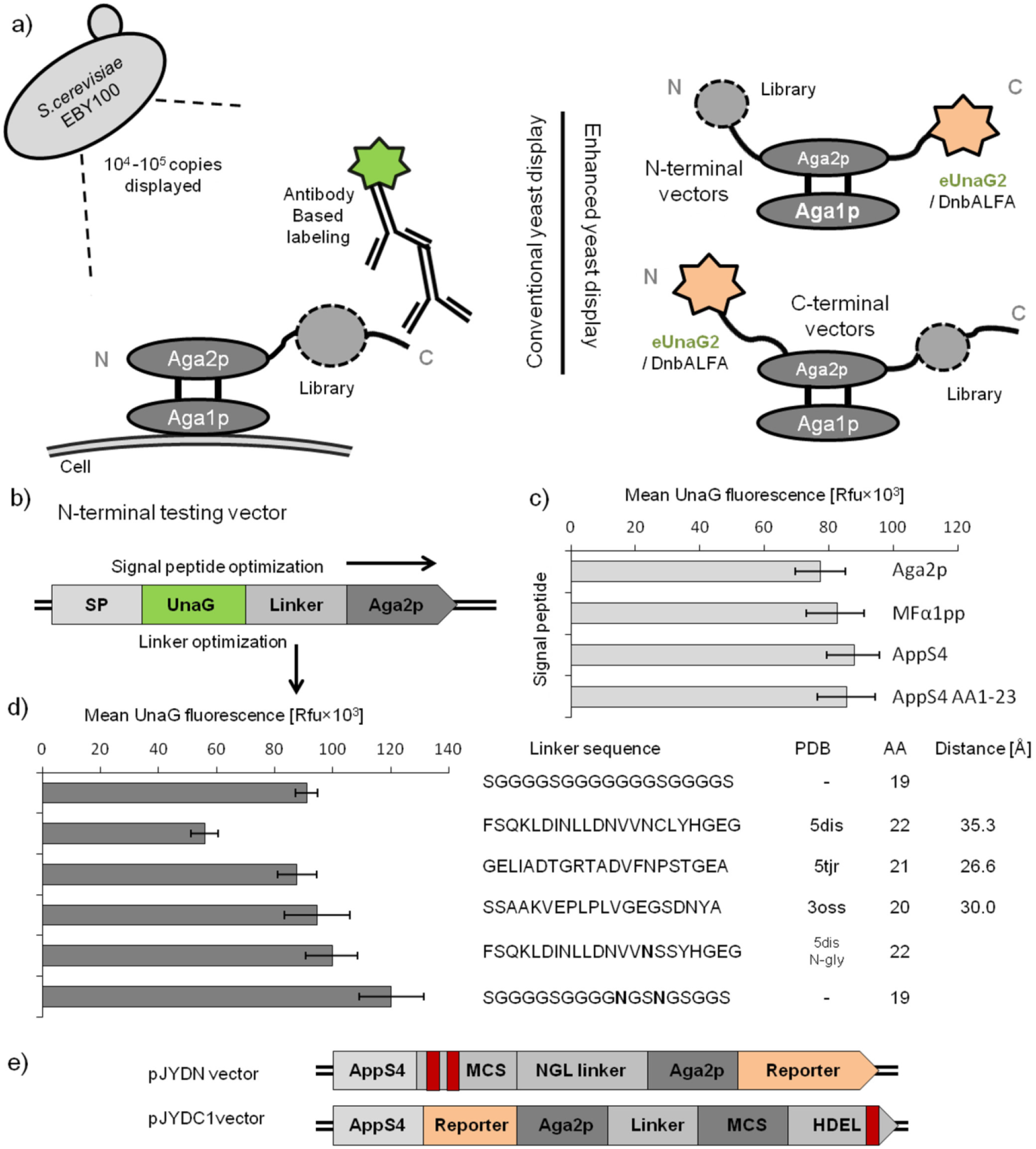
Schematic representation of enhanced yeast display constructs and their optimization. a) Schematic comparison of traditional yeast display based on mating agglutinin and enhanced yeast display. b) Expression unit organization for the signal peptide and linkers optimizations for N-terminal fusion vectors. c) Cytometry based comparison of the impact of different signal peptides on UnaG expression at the N-terminus of Aga2p. The mean fluorescence values were recorded for 30 000 yeast cells per sample. d) The impact of different linkers between UnaG and Aga2p proteins on fluorescence intensity assessed by flow cytometry measurements. The asparagine residues in linker sequences, highlighted in bold, were introduced in order to gain N-glycosylations. Except for glycine-serine stretches, flexible linkers were isolated from regions not resolved in electron density maps of corresponding structures {manuscript in preparation}. e) Final organization of N and C terminal vector. Stop codons are highlighted by red stripes.

To develop our plasmid system, we initially created a testing vector by introducing UnaG reporter between Aga2p native signal peptide and Aga2p (Figure 3b). Using this plasmid, we tested the impact of different signal peptides and linkers. We compared three different secretion signals: the natural Aga2p peptide, corresponding to AA 1 – 8, the engineered appS4 (Rakestraw et al., 2009), and the alpha mating factor 1 leader peptide (MFα1pp). The best performance was observed for the appS4 secretory leader (Figure 3c). The right peptide linker is important to prevent steric hindrance between the protein of interest and Aga2p and to optimize surface expression (Kjeldsen et al., 1997). We tested 4 different flexible linkers to secure a distance between yeast agglutinin and the displayed protein (Figure 3d). We tested the commonly used glycine – serine stretch (GGGGS)_x_ and 3 linkers developed in our lab. Linkers were mined from peptides invisible in protein crystallography structures {manuscript under preparation}. It is known that these sequences, not resolved in electron density maps, exhibit substantial flexibility (Schneider et al., 2014). The best linker was isolated from PDB 3oss. A free cysteine found in one of our linkers (PDB 5dis), which showed the lowest level of protein expression, motivated us to mutate this amino-acid residue. Instead of a simple change, we incorporate two serines instead of cysteine and its neighboring amino-acid leucine. This resulted in a gain of the N-glycosylation site in the linker sequence. These alterations resulted in a dramatic increase of eUnaG2 fluorescence (Figure 3d, second from bottom). In parallel, we tested the (GGGGS)_x_ linker with two asparagines incorporated, which resulted in the highest fluorescence, and was used for all further experiments (designated NGS linker, Figure 3d bottom). Overall, the linker engineering showed the importance of N-glycosylation for high yeast surface expression.

After signal peptide and linker optimization, the testing vector was rebuilt. Engineered reporters were incorporated at the C terminus between *Bam*HI and *Xho*I restriction sites and multiple cloning sites (MCS) were incorporated at 3 different positions in an adjacent region to the AppS4 leader peptide. Analysis of reporters expression showed the best position of MCS with the appS4 leader shortened (AA 1 – 23, Figure 3b). Plasmids containing eUnaG2 and DnbALFA were designed pJYDNp and pJYDN2p respectively. These plasmids do not contain stop codons, are expressed in appropriate galactose containing media and were used as expression controls.

Finally, on the basis of the optimized, new vectors, we created additional plasmids with two consecutive stop codons being introduced into the MCS (Figure 3e). This comes to avoid the possibility that the empty plasmid will give rise to a fluorescent signal. These plasmids are referred to as a negative plasmid (pJYDNn, pJYDN2n) and can be used either as negative control or template for plasmid cleavage and subsequent homologous recombination without the risk of false-positive colonies with empty plasmids.

#### Construction of C-terminal vectors

The C-terminal vectors resemble parental pCTcon2 organization with the protein of interest or library being fused to the C-terminus of Aga2p and reporters at the N-terminus of Aga2p (Figure 3a). This expression organization requires proper experimental design and controls because of the risk of a false-positive signal.

We used the above described N-terminal testing vector and cloned eUnaG2 at the N-terminus of the Aga2p and restored the original pCTcon2 MCS site at the C-terminus (Figure 3e top). The control experiment showed high eUnaG2 fluorescence in an empty vector. To limit this empty plasmid expression, we introduced *Saccharomyces cerevisiae* endoplasmic reticulum targeting peptide HDEL (Townsley et al., 1994) at the C-terminus of the MCS sequence between the *Nde*I and *BamH*I sites (Figure 3e bottom). Indeed, we confirmed using microscopy that the eUnaG2/DnbALFA-Aga2p-HDEL construct was predominantly retained in the endoplasmic reticulum (Figure S2a-c) with its fluorescence being reduced by almost 5-fold compared to the plasmid without retention signal. This is reducing the false positive signal from an empty plasmid emerging in library construction. To ensure that we detect only full-length constructs at the yeast surface, the C-terminal myc-tag was retained in the plasmid and we also created vectors with ALFA-tag to enable traditional labeling (pJYDC2). In addition, we confirmed protein retention in the endoplasmic reticulum for proteins hard to express – Uracil-DNA glycosylase (UniProt ID: P13051, Figure S2d) and used the combination of surface-specific labeling and eUnaG2 signal for visualization both intracellular protein and the cell surface-exposed fraction (Figure S2a-c). This approach allows for more controlled sorting in order to obtain clones with better cell surface exposure/secretion properties and to distinguish between proteins that are not secreted from those that are not expressed. We used the same approach to analyze that the HDEL escaping fraction which represents below 30 %.

#### Plasmid construction

Based on our optimized N and C terminal plasmids and engineered reporter proteins we constructed multiple yeast display vectors with different combinations of functional elements. The different plasmids and their functional element organizations are schematically shown in Table 1. All plasmids allow for labeling-free and traditional labeling procedures offering thus maximal versatility in the experimental design and monitoring of protein processing. A comparison between the pJYDNp plasmid expression (eUnaG2 reporter protein in yellow) and pJYDN2p plasmid expressing DnbALFA tagged mNeonGreen or miRFPnano670 is presented in the Figure 4b, showing the large gap achieved here between surface expressing yeasts to those that are not.

**Table 1.**
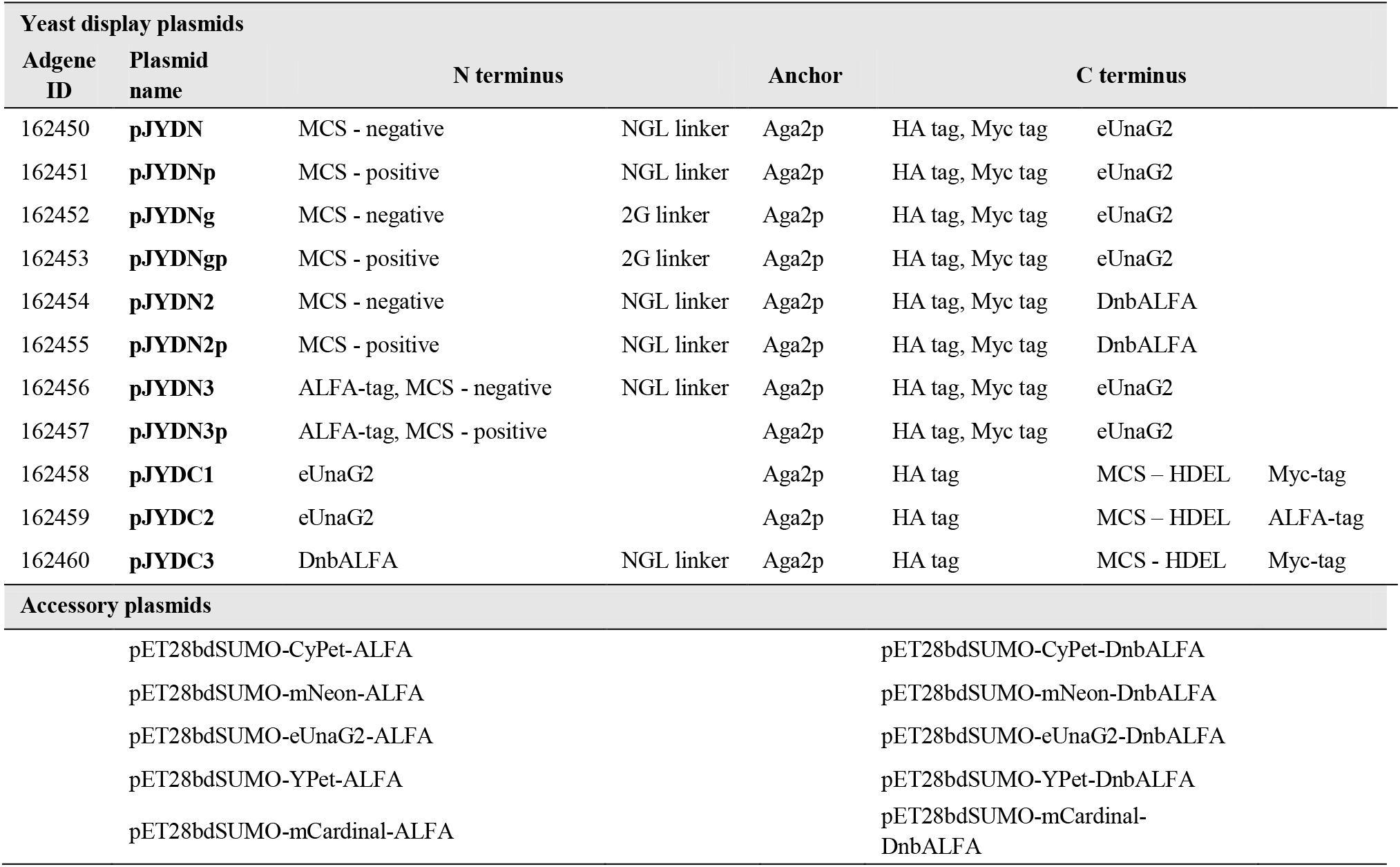
Summary of antibody labeling-free yeast display platform plasmids

**Figure 4.**
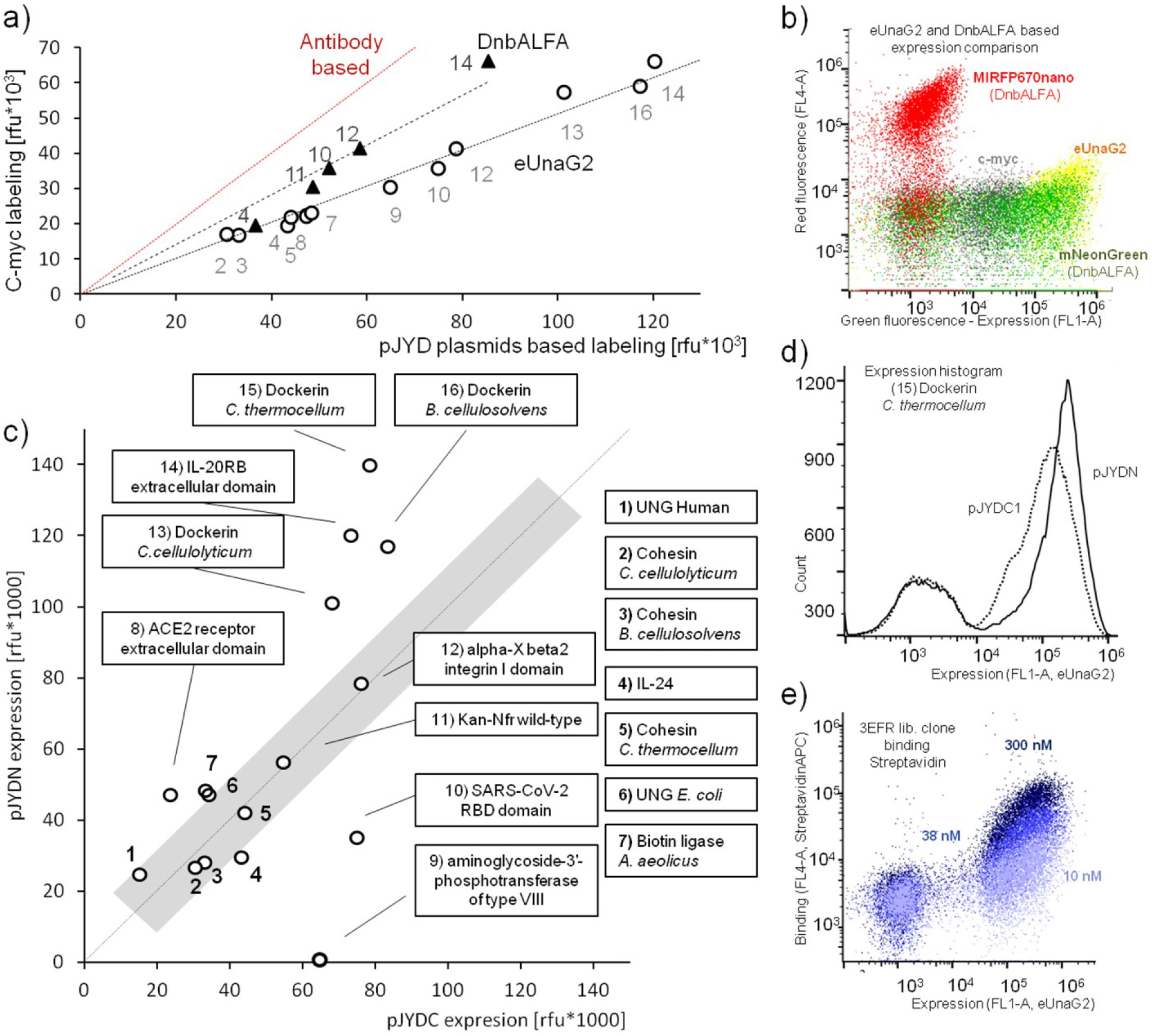
Comparison of expression between traditional and enhanced yeast display. a) Comparison of expression labeling intensities between traditional antibody-based c-myc labeling (pCTcon2) and the here engineered eUnaG2 (circles, pJYDC1) and DnbALFA (triangles, pJYDC3) alternatives. The numbers correspond to those in panel c, giving the identities of the proteins. b) FACS fluorescence dot plot signals comparison between eUnaG2 reporter (yellow), DnbALFA coupled with ALFA-mNeonGreen (green), or ALFA-miRFP670nano protein (red), and traditional anti-c-myc (grey). The eUnaG2 protein excitation maximum is at 498 nM and the emission maximum is 527 nM, which caused a small signal spillover into the red channel as evident at high signal intensities. A routine compensation procedure can be applied for signal correction. c) The differences in yeast surface expression between N (pJYDN) and C (pJYDC) terminal protein fusions with Aga2p among 16 tested proteins. The gray area highlights equal expression in both vectors (± 7,500 rfu). d) Overlay of expression histograms for dockerin from *C.thermocellum* (no. 15) expressed in pJYDC1 (C-terminal fusion with Aga2p) and pJYDN (N-terminal fusion). The comparison demonstrates higher expression and uniformity for dockerin fused with Aga2p at the N-terminus. e) Binding signal recorded together with eUnaG2 expression labeling. Stacked dot plots were acquired after incubation of 3EFR-Cfr-anti-StreptavidinAPC with 10, 38, and 300 nM of StreptavidinAPC for 1 h.

All yeast display plasmids were deposited in the Adgene plasmid repository and their corresponding ID numbers are shown in Table 1. Table 1 also shows the plasmids constructed to complement our yeast display vectors with vectors for the production of fluorescent proteins: ALFA-tagged fluorescent proteins and fluorescent proteins fused with DnbALFA. The expression vectors are based on pET28bdSUMO vector (Zahradnik et al., 2019) and enable for bdSUMO protease single step, on-column cleavage based, purifications (Frey and Görlich, 2014). An example of proteins purified by this single step purification process is shown in Figure S3. Both ALFA-tagged proteins and DnbALFA fusions do not require further purification steps and can be used directly for co-cultivation labeling.

#### Examining protein expression using the enhanced yeast display platform

To test the applicability of our yeast display system, we analyzed the expression of 16 proteins by using traditional anti-c-myc antibody-based labeling with secondary antibody conjugated to Alexa Fluor 488 (Chao et al., 2006)(pCTcon2 plasmid), intrinsic eUnaG2 fluorescent signal (pJYDC1 plasmid), and DnbALFA with co-cultivation labeling with Alfa-mNeonGreen (pJYDC3 plasmid). Our results show a very tight correlation in the strength of the fluorescence signal between the three systems. This suggests that the expression level of the proteins is similar using either platform. However, the absolute fluorescence intensities were the brightest for eUnaG2 (double of c-myc) followed by DnbALFA – mNeonGreen labeling (50% increase over c-myc), Figure 4a and b.

Next, we tested the differences in expression levels between the two basic plasmid arrangements: the protein being N or C terminal to Aga2p. As a fluorescence probe, we used eUnaG2 fused C and N terminal to Aga2p in plasmids pJYDN and pJYDC1 respectively (Figure 4c). Large variations in levels of expression were identified among the tested proteins. The dockerin proteins (*Clostridium cellulolyticum*, sequence ID: M93096.1; *Bacteroides cellulosolvens,* AF224509.3; *Clostridium thermocellum*, L06942.1), the receptor IL-20RB, and the angiotensin-converting enzyme ACE2 are preferentially expressed as N-terminal fusions, in contrast to kanamycin resistance protein (Boyko et al., 2016) and biotin ligase ID2 from *Aquifex aeolicus* (Tron et al., 2009) which best express as C-terminal fusions. Our inspection of 3D structures for all tested proteins explained only the lack of expression for the kanamycin resistance protein PDB ID 4H05. The strictly conserved C-terminus of this protein was found buried deep inside the structure, not allowing for C-terminus modifications. Therefore, we suggest experimental expression testing for every construct and its expression optimization. Still, if possible, N-terminal expression is preferable as it purges stop-codon insertions in the library without the need for additional steps.

### Part II: evaluating the efficiency of the enhanced yeast display platform for the emergence of new protein-protein interactions between random proteins

#### Protein libraries construction

The here created pJYDNn and pJYDNg plasmids were used for the generation of 6 targeted saturation mutagenesis protein libraries (Table 2). The proteins include 4 previously published scaffold proteins and two new candidates – aminoglycoside-3’-phosphotransferase of type VIII N-terminal domain fragment (Kan-Nfr) and biotin ligase ID2 from *Aquifex aeolicus* C-terminal domain fragment (3EFR-Cfr). Both new candidates were chosen to test the possibility of *in vivo* enzyme complementation-based experiments that are not covered within the scope of this publication. Among the proteins, Sso7d, Knottin, and GP2 have very high melting temperature. The other three were pre-stabilized before library preparation using PROSS calculations and subsequent selection of the suggested mutations for the highest level of expression on yeast surface, using yeast display. This resulted in the incorporation of 5 stabilizing mutations in s3LYV, 7 mutations in 3EFR-Cfr, and 13 mutations in Kan-Nfr (Supplementary Material S1 and Figure S4). The corresponding change in melting temperature upon stabilization was measured using the Prometheus NT.48 for the 3EFR-Cfr and 4H05-Nfr purified wild type and designed proteins. The 3EFR-Cfr, and 4H05-Nfr stabilized protein variants were 11°C and >20°C more stable than the starting proteins. The s3LYV stabilized protein showed almost 20 % higher expression on the yeast surface and better expression in *E.coli* with reduction of inclusion bodies formation.

**Table 2.**
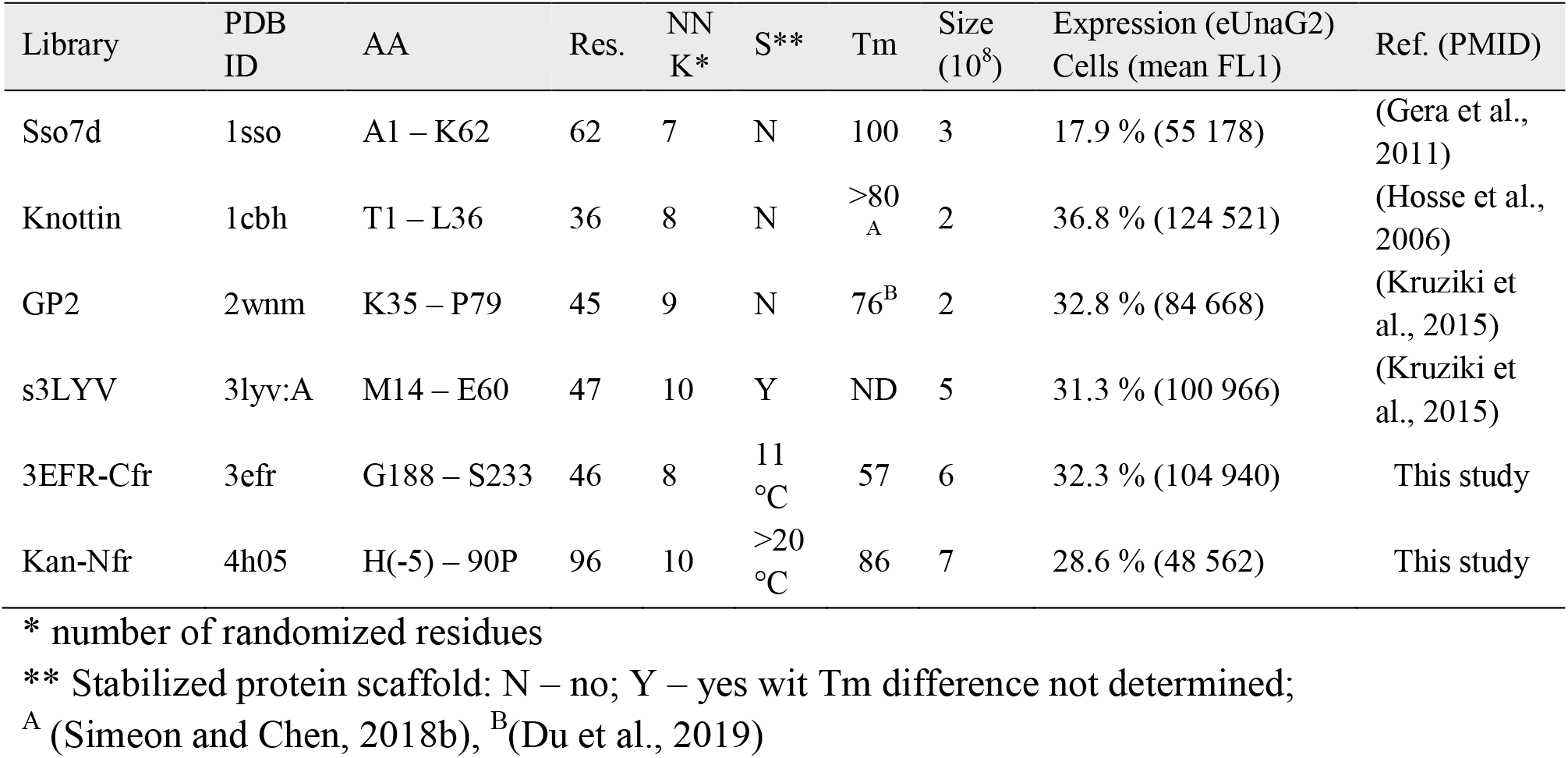
pJYDNn plasmid-based yeast display libraries

For library construction we chose to randomize specific, structurally clustered positions, providing coverage of all possible mutations and combinations rather than random mutagenesis of the complete protein. For Kan-Nfr, positions for library construction were identified by a combination of multiple sequence alignment and *in silico* FoldEX (Schymkowitz et al., 2005) based saturation mutagenesis (Figure S5). Positions and patches in the structure with large evolutionary variability but low energy variability between mutations were targeted. For 3EFR-Cfr, the small scaffold size made us randomize the β-sheets connecting loops. The 6 tested scaffolds are comparable in sequence length - all are very small proteins. The outcoming library sizes were comparable, with the number of randomized positions being 7 – 10 (Table 2). The structure, sequence, and exact position of randomized residues are shown in Supplementary Material S2. Library qualities were verified by sequencing 20 randomly selected clones.

#### High stringency selection for tight binders

The first selection was aimed to find high-affinity binding variants to the commercially available Streptavidin-APC conjugate protein as bait. Our selection strategy was based on pre-selection against high concentration of Streptavidin-APC, to decrease the complexity of libraries, and subsequent construction of a new pooled library of all pre-selected scaffold variants (Figure 5a left). In the first round, we used all our naïve libraries independently against 1 μM target protein and sorted approximately 1 % of cells in the binding/expression quadrant (double-positive cells). In total, we sorted slightly above 106 yeast cells from each library. In the second step, all selected clones from the different libraries were pooled, while keeping the same number of clones from each library (10^7^ per library). Subsequent rounds of sorting were done with the pooled library against decreasing concentrations of the bait protein – 500 nM, 100 nM, 50 nM, and finally 25 nM Streptavidin-APC. The population of the last sort was plated and 20 single colonies were screened for binding and sequencing. Among all sequences, we identified one dominant (19/20) 3EFR-Cfr library clone. The equilibrium dissociation constant measured by flow-cytometry was calculated to be 28 ± 1.6 nM. The other, single clone was a member of the s3LYV library and its affinity was estimated to be ≥ 500 nM (Figure 5b).

**Figure 5.**
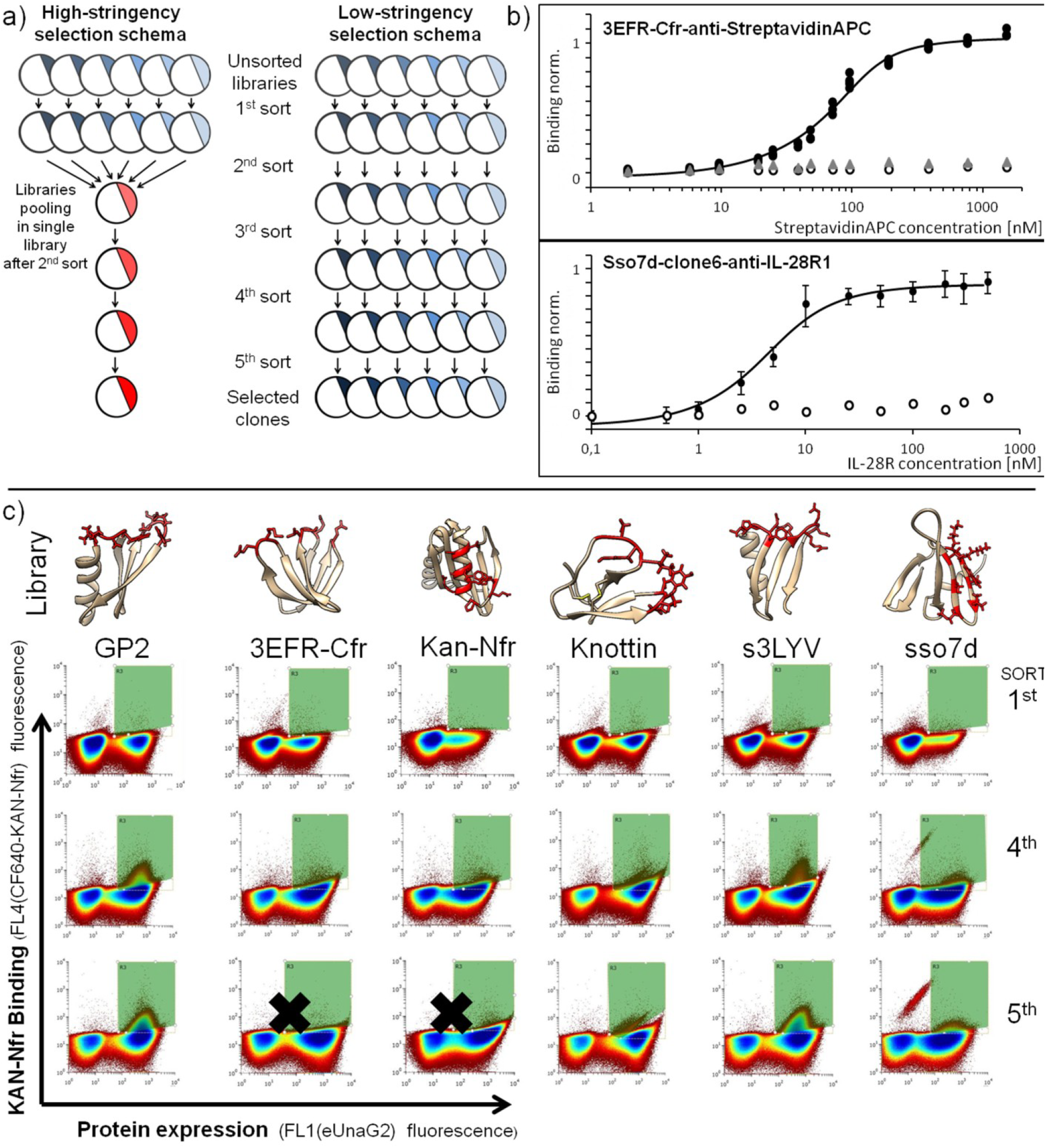
Selections and emergence of new protein-protein interactions between random proteins using enhanced yeast display. a) Schema of high (left) and low (right) stringency selections used with all six scaffold libraries. b) Binding of selected clones identified after high stringency selection. High-affinity binder – black circle data points (triplicates); empty circles – wild type scaffold; black triangles – additional clones; grey triangles – 3EFR-Cfr wild type (non-stabilized) with introduced StreptavidinAPC binding residues; c) Low stringency selection process overview among all used libraries. The red amino-acid residues in the structure representation illustrate the randomized positions. Each FACS-sorting dot plot represents 10^6^ events. The green area is the sorted region. Sso7d library plots were analyzed together with a sorting control – red fluorescent protein mCardinal expressing EBY100 cells. Libraries with non-significant enrichment are marked.

To demonstrate the importance of pre-stabilization of 3EFR-Cfr prior to selection, we transferred the 3EFR-Cfr-Anti-StreptavidinAPC binding residues to the non-stabilized 3EFR-Cfr and tested its expression and binding properties. The comparison between stabilized and wild type scaffolds showed complete loss of StreptavidinAPC binding on the wild type scaffold, and a reduction of the clone’s expression by 14 % (Figure 5b).

Using the same prey libraries, we now repeated the selection against the purified extracellular portion of IL-28R1, the high-affinity receptor for interferon lambda, as bait. In the first round, all libraries were selected independently against 1 μM protein, and then they were pooled and subjected for additional rounds of FACS selection with decreasing concentration of bait – 500, 200, 100, and 50 mM. After the selection, 20 colonies were isolated and screened for binding and sequencing. Two different sso7d clones were identified. The most prevalent sso7d clone (no. 6, 19/20 sequences) had a binding affinity of 2.4 ± 1.1 nM as measured by cytometry binding analysis (Figure 5b). The second clone (no. 9) had a *K*_D_ > 1 μM, which corresponds to its incidental presence among sequenced clones.

#### Low-stringency selection to identify sequence space allowing for weak binding

The low-stringency selection experiments were done using the purified Kan-Nfr protein fragment as bait. The selection strategy was set to demonstrate the selections of independent libraries to a specific protein at a given concentration. No pooled library was constructed (Figure 5a, right). In the first round, we selected naïve libraries against 5 μM Kan-Nfr protein and sorted approximately 0.1 % of cells in the binding/expression quadrant (double-positive cells). A minimum of 20 000 cells was sorted in each library and selection step. The sorting schema was kept the same with concentrations as follows: 5, 2.5, 1, and 0.5 μM. The representative dot plots from library sorting are depicted in Figure 5c.

For all 6 libraries, one observes much higher fractions of yeast cells expressing the target protein between sort 1 and 4, due to the positive selection against eUnaG2. The 3EFR-Cfr and Kan-Nfr libraries did not show any enrichment for Kan-Nfr binding during the selection rounds. Knottin and sso7d libraries showed enrichment of 1 and 3 % respectively during the last selection round and 500 nM bait protein. Both libraries showed up to a 20% increase in red fluorescence only. This suggests that low-affinity binding clones are dominant in these libraries. The highest enrichment reaching up to 10 % was recorded in GP2 and s3LYV libraries. Both libraries also showed an order of magnitude difference in fluorescence levels compare to non-binding cells.

We randomly selected 20 colonies from the four libraries where enrichment is seen and sequenced them. The 20 selected clones demonstrated considerable variability among the sequenced clones. We identified 11 different sequences within the GP2 library selected clones (Figure 6a). All of the single clones were tested for binding against 500 nM of Kan-Nfr, with full binding curves being determined for the best two binding proteins: GP2-G1 and G2. These two proteins, with affinities of 1 μM and 0.83 μM have considerably higher affinity than the other candidates, which have estimated affinities higher than 1.5 μM (Figure 6b). Rapid inspection of the structure of wild-type GP2 (PDB ID 2WNM) supported the formation of a new disulfide bond in the G1 clone (Figure 6a). Other clones were clustered in three different groups of sequences by Maximum Likelihood phylogenic analysis (Figure S6). Members of each group shared some amino-acid residues or chemical properties suggesting enrichment for multiple binding modes or interfaces among clones. Screening of the s3LYV sorted library yielded 12 unique sequences. Their sequencing showed a similar picture to that obtained for the GP2 library, yet with lower affinities. One clone bound with an affinity of ~3.5 μM, with the other clones having affinities between 3.5 and 7.0 μM (Figure S7).

**Figure 6.**
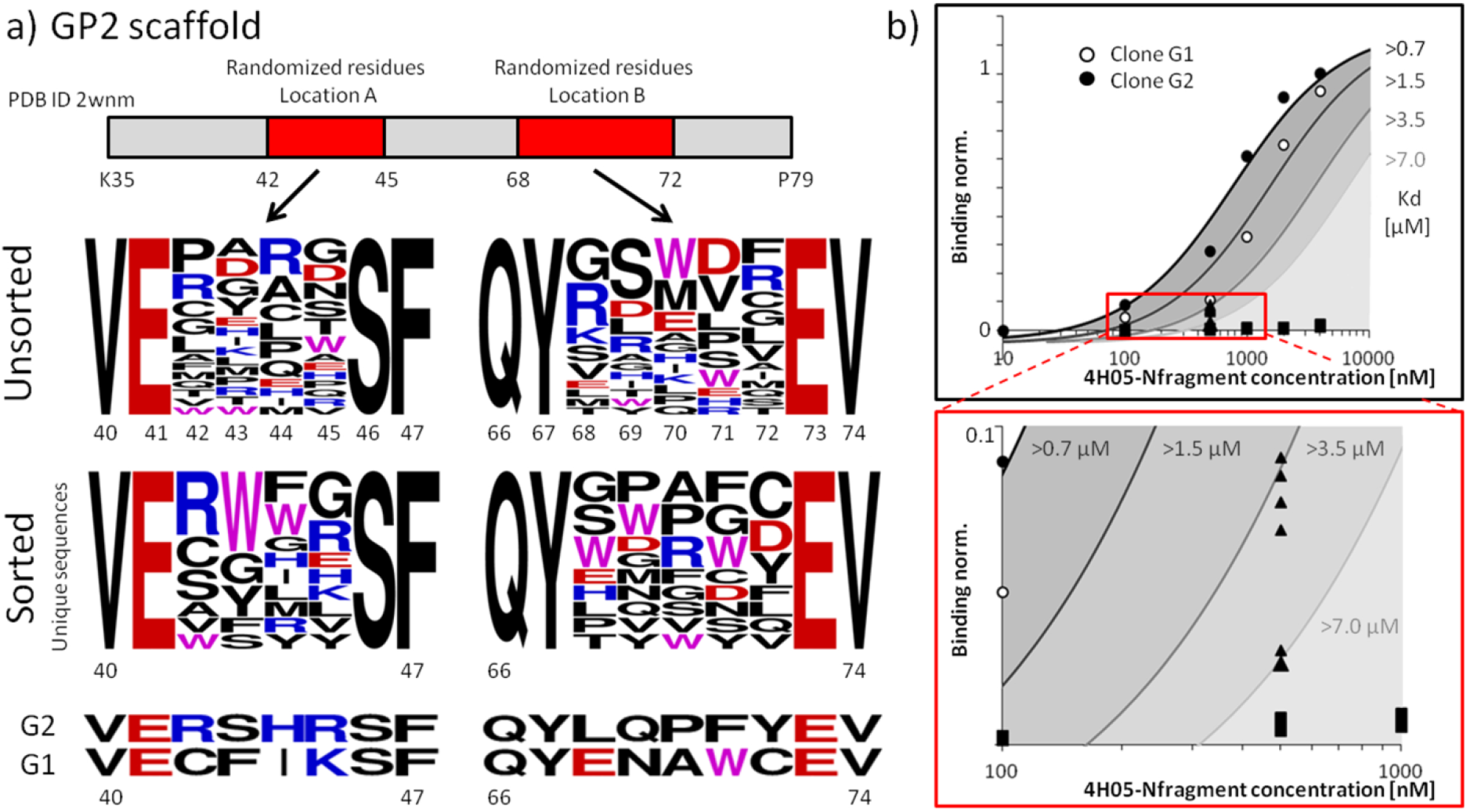
Low stringency selection of the GP2 scaffold library against Kan-Nfr. a) Comparison of non-redundant sequences of randomly selected clones from the unsorted and sorted GP2 library. Six clones from the unsorted library contained stop codons, which is not reflected in the sequence logo. The logo was created by using Sequence Logo Generator (Crooks et al., 2004) and the randomized residues with two non-randomized amino acid residues from each side. b) Measuring binding affinities for the 11 screened clones. The binding curves were determined for G1 and G2 clones (circles). The fitted curves were used to estimate binding affinities for clones with weaker binding (black triangles). The wild type background values are depicted by black squares (triplicates). The lower plot is a zoom-out of the red area in the upper plot.

The Knottin and sso7d selected libraries showed much lower fluorescence intensities during enrichment towards Kan-Nfr than the s3LYV and GP2 libraries. The analysis of the Knottin library identified only 3 different clones with considerable sequence similarities (Figure S7). We hypothesize that this corresponds to a low level of library enrichment and consistently low number of binding clones. The binding assay with 500 nM bait showed affinities higher than 7 μM, which corresponds to a low fluorescence increase in the binding population during the sorting process. The sso7d library screening yielded 6 different clones also with some sequence similarities resembling the situation in the Knottin library. The binding affinity for sso7d library clones can be estimated even lower than for knottin clones with values ≥ 10 μM. We did not further investigate these two libraries.

## Discussion

Yeast display is maybe the most successful of selection methods, with a wide array of applications. Still, labeling of yeast cells for flow cytometry analysis is laborious and cost-ineffective. Its requirements for multiple washing steps and long incubation times with antibodies (Chao et al., 2006) motivated researchers to search for process simplifications. The use of GFP fluorescent protein (Lim et al., 2017; Ye et al., 2000) and ACP, an orthogonal acyl carrier protein (Uchański et al., 2019) for simplification of yeast surface display was reported. ACP shows excellent labeling properties but CoA-biotin and fluorescent CoA-547, CoA-647 derivatives are needed as well as 1 h of incubation. The use of GFP in the secretory pathway is connected with multiple impediments such as protein targeting to the vacuole (Kunze et al., 1999; Li et al., 2002). Some of these difficulties were solved by protein engineering (Eiden-Plach et al., 2004; Huang and Shusta, 2005b; Roh et al., 2013;Grzeschik et al., 2017). Other problems, like the inability to turn off the fluorescence of GFP, cannot be solved. Overall, none of the alternative techniques became dominant, replacing the method published by Chao (Chao et al., 2006).

Since we aim to use yeast display to investigate the potential evolution of new protein-protein interactions, and how they can emerge for randomly chosen proteins, we decided to first optimize yeast display, allowing for multiple expression and labeling strategies. The first step was to rebuild the most commonly used vector for yeast display – pCTcon2 by the extensive trimming of all the unnecessary plasmid parts. The resulting vector, pJYD, is ~20 % smaller than the parental vector, yet keeping all the critical parts and showing similar properties. Based on this new plasmid backbone we tested the suitability of several reporter proteins: four green fluorescent proteins, five far-red fluorescent proteins, and two peptide tag-recognizing nanobodies. All proteins were expressed, under the control of the galactose-GAL promoter, both in the cell cytoplasm and on the yeast surface tethered to Aga2p yeast mating agglutinin. The differences between the cytoplasmic and surface expressions suggested the protein’s effectiveness in being exposed on the yeast surface. Considering the differences in expression, protein size, and brightness we identified UnaG, a bilirubin dependent fluorescent protein, and ALFA-tag binding nanobody as the best candidates for protein engineering. Both reporters enabled regulation of their signal and change in the labeling strategy upon need.

Reporter proteins UnaG and nbALFA were subjected to multiple rounds of protein engineering to tailor their properties to fit the yeast surface display platform. Proteins were stabilized by using a combination of PROSS calculations (Goldenzweig et al., 2016), FACS selections, and N-linked glycosylation sites introduction. In total, we introduced 10 mutations in UnaG, with the optimized variant being called eUnaG2 (Figure 2). From nbALFA, we created the DnbALFA protein, which differs by 10 mutations. The eUnag2 average fluorescence intensity in the expressing population was two and five-fold higher compared to eUnaG (Yeh et al., 2017) and UnaG respectively (Kumagai et al., 2013). The affinity of eUnaG2 for bilirubin was measured and is slightly higher (46 ±13 pM, Figure S1) than 98 pM reported for UnaG (Kumagai et al., 2013). The expression of DnbALFA is almost 5 times better in mean fluorescence values than the expression of nbALFA on the surface of *S. cerevisiae* EBY100 cells. Interestingly, the protein expression in *E. coli* BL21 (DE3) was also highly enhanced, despite the lack of glycosylation, which resulted in lower melting temperature compared to wild type. The SDS-PAGE expression analysis showed more than 10-times higher yield for the designed protein variant (Figure S1).

The tailored reporter proteins, eUnaG2 and DnbALFA, were used as the basis for the construction of a whole vector platform allowing for different expression construct organizations and multiple detection options. The construction of the N-terminal Aga2p fusion vector was accompanied by testing of suitable linkers and signal peptide sequences to optimize the vector properties. An impact of signal peptides on expression has been shown previously (Rakestraw et al., 2009). Vectors for the C terminal fusions to Aga2p protein were modified with endoplasmic retention signal to reduce the possibility of false-positive signals (Figure 3). Interestingly, the HDEL retention signal control is not perfect and there is still an escaping population (Figure S2). This might be caused by high protein load in the endoplasmic reticulum or by the pulling effect caused by fusion with Aga1p. We recommend careful control experiment or C-terminal tag labeling in the first round of selection.

To confirm the applicability of our vectors and to exclude the possibility of the reporter protein influencing expression, we compared protein surface expression of 16 different proteins fused to the C-terminus of Aga2p in pCTcon2 and pJYDC (Figure 4). Protein expression was assessed both by c-myc labeling and by intrinsic fluorescence of eUnaG2 (pJYDC). eUnaG2 fusion resulted in double the fluorescence in comparison with c-myc for all the proteins tested. Besides this comparison, we analyzed the variation in expression of N and C terminal fusions. Most of our proteins were expressed in both N and C terminal vectors with an almost similar level of expression. 5 proteins were preferentially expressed as N terminal fusions (3 dockerins, IL-20RB and ACE2), while SARS-CoV-2 RBD and the 4H05 proteins were preferentially expressed as C terminal fusions. Overall, these experiments show that if possible, N-terminal insertion of the library, using eUnaG2 as fluorescence marker for expression is the most desirable method (as it purges stop-codons from the library), however, if needed C-terminal fusion, as well as the use of the far red-fluorescence protein miRFP670nano fused to DnbALFA also gives excellent results and ease of use.

Having created an enhanced, high-throughput yeast display platform, we tested our ability to use it for evolving new protein-protein binding sites. First, we generated 6 targeted protein libraries using the pJYDNn plasmid for six different scaffold proteins. Among them, 4 scaffolds were based on literature published data, and 2 scaffolds were developed by us (Table 2, Figure S4, S5). Stabilization design was implemented for 3 scaffolds before library construction (s3LYV, 3EFR-Cfr, and Kan-Nfr; Supplementary Material S1, S2), which has been shown to have a dramatic impact on proteins’ evolvability (Bloom et al., 2006; Tokuriki and Tawfik, 2009) by expanding their mutational space. The melting temperature of other scaffolds used in this study was already high (Table 2). The libraries differed in the number of randomized residues and the estimated complexity depending on the design and yeast homologous recombination quality (Table 2).

We used two fundamentally different selection strategies to demonstrate the likelihood for binding (Figure 5a). Initially, we used stringent conditions and a pooled library after an initial selection round. This experimental strategy allows for different scaffolds to compete with each other in the selection process and selection of the best clones among them. We identified high-affinity binders towards the two baits, streptavidin (28 ± 1.6 nM, 3EFR library clone, Figure 5b) and IL-28R1 (2.4 ± 1.1 nM, sso7d library clone, Figure 5b). These values are comparable with binders obtained by methods using much higher complexity libraries (Wilson et al., 2001). The affinity for IL-28R1 was even higher than the binding affinity with its natural ligands as measured by ELISA assay (15 nM for IFNL1 and 65 nM for IFNL3) (Au - Syedbasha et al., 2016). By applying stringent conditions, we identified only a single high-affinity clone for each of the two baits, among the 20 sequenced colonies. This indicates that the best clone over-competes others during selection cycles. Both high-affinity binders did not originate from the most complex library. Moreover, the sso7d library had the lowest proportion of expressing cells, yet gave rise to the best binder to IL-28R1. It demonstrates the importance of parallelization with different libraries in order to select for high-affinity binders since both chemical and shape complementarity are important for binding (Schreiber and Fleishman, 2013). The fact that we obtained high-affinity binders for both target proteins, done within a week’s time, shows the power of our approach. This would suggest, that using the pre-made libraries of these 6 extremely small proteins is sufficient to fish for high-affinity binders for a large variety of proteins. In addition, the results show the importance of using highly stable proteins to increase the success of library selection, as the control experiment where we introduced the wild type residues back to the StreptavidinAPC binding 3EFR clone showed a complete loss of binding affinity.

In previous research, we demonstrated the high plasticity of TEM1 – BLIP interface under low-stringency selection (Cohen-Khait and Schreiber, 2016) and the limited number of mutations needed for self-interaction (Cohen-Khait et al., 2017). Results suggest that multiple solutions for binding exist and there is only a limited barrier between binding and nonbinding proteins. Here, we used the low-stringency selection strategy to further validate those data. Using μM quantities of the target Kan-Nfr protein, we were able to select binding clones from the Knottin, GP2, s3LYV, and sso7d libraries, showing enrichment in the expressing and binding populations (Figure 5c), and subsequently multiple binding clones with affinities in the μM range (Figure 6). Moreover, multiple low-affinity solutions were identified. This implicates that low-affinity binding can be easily evolved, basically between almost any protein surface with a second protein. *In vitro*, this can be achieved with almost every bait pair library by the acquisition of a low number of mutations. This result supports the hypothesis that restricting low-affinity interactions between non-cognate pairs is an important evolutionary force dictating properties of the protein surfaces especially in abundant proteins and highly influencing the complexity of proteomes (Agozzino and Dill, 2018; Yang et al., 2012).

## Conclusions

We applied a combination of restriction-free cloning methods, plasmid optimizations, and protein engineering to enhance the widely used yeast display method and simplify the current methodological approach. We constructed 11 different yeast display vectors to enable the N-terminal, C-terminal fusions, and multiple labeling options. The created plasmid platform outperforms traditional antibody-based methods by reduction of hand on time, cost-effectiveness, and higher accuracy with both low and very high binding affinities. Thanks to these advantages, it enables sorting parallelization without the need for robotized FACS screening platforms.

We evaluated the enhanced platform on 6 different protein libraries to uncover the differences in selection under high and low stringency selection conditions. High-affinity binders were selected from a single library which dominated selection after 5 rounds, suggesting that the high-affinity clone outcompetes the others. In contrast to the high-stringency conditions, the selection under low-stringency conditions led to the identification of multiple binding clones in the majority of libraries, pointing towards the many solutions available for low-affinity binding. These experiments lead to two main conclusions. Firstly, the parallelization leads to an optimal scaffold selection and isolation of high-affinity binders without the need for high complexity libraries to be synthesized. Secondly, to create a weak binding *de novo* is an easy task to achieve and probably there is an evolutionary selection against it.

Besides the above-mentioned conclusions, we described two new scaffold proteins for the selection of high-affinity binders that were created by stabilization of protein fragments and the application of restriction-free cloning for library preparation. Both approaches are helping to simplify the overall process of design of making libraries. Our work should help to expand the range of uses for yeast display and could support discoveries by using this exciting method.

## Materials and methods

### pJYD yeast display backbone construction and DNA manipulations

The pJYD vector backbone was assembled by the three-components assembly (Peleg and Unger, 2014) from pCTcon2_KAN_ vector (Cohen-Khait and Schreiber, 2016). All components were PCR amplified by using KAPA HiFi HotStart ReadyMix (Roche, Switzerland), the template vector was removed by *Dpn*I treatment (NEB, USA) at 37°C (1 – 2 h) and subsequently, the amplicons were purified by NucleoSpin® Gel and PCR Clean-up kit (Nachery-Nagel, Germany). The assembly reaction was composed of 100 ng of each amplicon and KAPA HiFi HotStart ReadyMix (50 ul reaction mix). The reaction was divided into 5 aliquots (10 ul) and subjected to assembly PCR (30 cycles; 1 min annealing; 60 – 70°C gradient; with 2°C increments per aliquot; 6 min of polymerization). 1 μl from PCR reaction aliquots were transformed in electrocompetent *E.coli* Cloni® 10G cells (Lucigen, USA). Colonies were screened by colony PCR and positive colonies were sequenced. The whole plasmid sequence was verified.

Incorporation of further changes in pJYD vectors and other cloning was done via the restriction-free cloning procedure (van den Ent and Löwe, 2006). The mutagenic primers were used for amplification of megaprimers. If incorporation or modification of long sequence was needed, multiple extension PCR amplifications were applied with overlapping primers. All PCR reactions were done by using KAPA HiFi HotStart ReadyMix (Roche, Switzerland). Purified megaprimers (200 ng of DNA, Nachery-Nagel, Germany) were mixed with 20 ng of destination plasmid and subjected to PCR similar to assembly reaction. The template vector was removed from the PCR mixture by *Dpn*I treatment (NEB, USA) at 37°C (1 – 2 h) and 1 μl from PCR reaction aliquots were transformed in electrocompetent *E.coli* Cloni® 10G cells (Lucigen, USA). Colonies were screened by colony PCR and sequenced.

### Reporter genes and protein engineering

DNA fragments of mNeonGreen (Shaner et al., 2013), yeGFP (Huang and Shusta, 2005a), UnaG (Kumagai et al., 2013), iLOV (Chapman et al., 2008), dFP-mini (Sheehan et al., 2018), GAF-FP (Rumyantsev et al., 2015), TDsmURFP (Rodriguez et al., 2016), IFP1.4 (Shcherbakova and Verkhusha, 2013), miRFP670nano (Oliinyk et al., 2019), nbBC2 (Braun et al., 2016), and nbALFA (Götzke et al., 2019) were ordered from Twist Bioscience (USA) with *S. cerevisiae* codon optimization. Reporter genes were amplified by KAPA HiFi HotStart ReadyMix (Roche, Switzerland) with two sets of primers for intracellular expression (starting with ATG and omitting the Aga2p secretion signal) and yeast surface expression (insertion between *Bam*HI and *Bgl*II sites). PCR products were purified by using NucleoSpin® Gel and PCR Clean-up kit (Nachery-Nagel, Germany) and used for amplicon incorporation in destination plasmid by subsequent PCR. The template plasmid molecules were inactivated by *Dpn*I treatment (1 h, NEB, USA) and directly transformed to electrocompetent *E. coli* Cloni® 10G cells (Lucigen, USA; 1ul crude reaction mix). Kanamycin selected clones were screened by colony PCR and verified by sequencing. Correct plasmids were transformed in the EBY100 *Saccharomyces cerevisiae* strain by the lithium acetate method (Gietz and Woods, 2002) and grown on yeast minimal SD-W plates. Reporters’ expressions were analyzed for five colonies by bdAccuri cytometer (BD Life Sciences, USA).

Proteins UnaG (Kumagai et al., 2013), nbALFA (Götzke et al., 2019), and miRFP670nano (Oliinyk et al., 2019) were subjected for prediction of stabilizing mutations in Pross server (Goldenzweig et al., 2016). Mutagenic primers with suggested mutations were used to generate random libraries via restriction-free method TPCR described by Erijman et. al (Erijman et al., 2011). All colonies grown on selection plates were pooled, subjected for mini-prep plasmid purification by Wizard® Plus Minipreps DNA Purification System (Promega, USA), and used as the template in subsequent PCR amplification of the given library. The amplicons were purified (NucleoSpin®, Nachery-Nagel, Germany), mixed with the purified cleaved plasmid (pCTcon2KAN vector (Cohen-Khait and Schreiber, 2016), NheI and BglII), and used for yeast transformation (Benatuil et al., 2010). Yeast cell cultivation, expression, and selection procedures are described in specific chapters. Cells accompanied with higher fluorescence intensities were sorted, cultivated, and their plasmids were isolated. Isolated plasmids were sequenced and mutations were analyzed (20 colonies). New genes, including all needed mutations, were purchased from Twist Bioscience (USA) and additional modifications such as gain of N-glycosylation were introduced by site-directed mutagenesis using restriction-free cloning (Peleg and Unger, 2014).

### DNA libraries preparation

All libraries were constructed by consecutive extension PCR amplification by using KAPA HiFi HotStart ReadyMix (Roche, Switzerland) and NNK randomized primers (Sigma, USA). The 3EFR-Cfr and GP2 libraries were constructed by extension amplifications of Aga2p gene together with NGL linker. The Knottin library was constructed by the same approach with pJYDNg plasmid. No template genes were used for the construction of these libraries. In contrast, the Kan-Nfr, sso7d, and s3LYV libraries were amplified from template DNA (pET28bdSUMO plasmids (Zahradnik et al., 2019)). In order to reduce the possibility of template amplification, the DNA was gel purified between each PCR extension step. Alterations in codon usage were incorporated in primers to further reduce the template amplification possibility. Library sequences are shown in Supplementary material S13 and the corresponding primers are shown in the Supplementary material S16. Purified DNA (10 – 20 μg per library) was mixed with NdeI and BamHI cleaved pJYDNn or pJYDNg plasmids (4 μg) and electroporated to EBY100 *Saccharomyces cerevisiae* (Benatuil et al., 2010).

### Recombinant protein expression systems and purification

The extracellular part of IL-28R1 (UniProt ID <u>Q8IU57</u>, AA 21-228) was produced by the *Drosophila* S2 expression system. The gene optimized for the *Drosophila* codon usage and extended by C-terminal His-tag was purchased from Life Technologies (DNA String fragments, USA). The DNA fragment was inserted into a pMT-BiP-V5-His_A vector (ThermoFisher) by using restriction-free cloning (Peleg and Unger, 2014) between the restriction sites *Bgl*II and *Xho*I and verified. Purified plasmid (Plasmid Plus Midi Kit, QIAGEN, Germany) was mixed with selection plasmid pCoBlast (1:10) and the mixture was used for cells transfection by Effectene Transfection Reagent (QIAGEN, Germany) according to the manufacturer’s protocol. A stable cell line was selected by using 25 μg/mL Blasticidin S and protein expression was induced by 1.0 mM CuSO4. The protein purification from precipitated cell-culture media was done on HisTrap HP 5 ml and HiLoad 16/600 Superdex 75 (GE Healthcare, USA) columns by the method described previously (Zahradník et al., 2018).

Protein expressions based on pET26b and pET28bdSUMO (Zahradnik et al., 2019) were done by using *E. coli* BL21(DE3) cells and 200 ml 2YT media (1L Erlenmeyer flasks). Cell cultures were grown (30°C, 220 rpm) to OD600 = 0.6, then the temperature was lowered to 20°C, protein expression was induced by 0.5 mM IPTG, and growth continued for 12 – 16h. Cells were harvested (6000g, 10 min), disintegrated by sonication in 50 mM Tris-HCl, 200 mM NaCl buffer (pH 8), and purified by NiNTA agarose. Proteins fused with SUMO (pET28bdSUMO plasmid) were purified by the on-column cleavage method (Frey and Görlich, 2014). Eluted fractions were analyzed on SDS-PAGE gels. For higher purity HiLoad 26/600 Superdex 75 gel filtration chromatography was applied (PBS buffer).

### Yeast transformation, cultivation, and expression procedures

pJYD plasmids were transformed into *S. cerevisiae* EBY100 by LiAc – PEG method (Gietz and Woods, 2002) and grown on yeast minimal SD-W plates 48 – 72 h at 30°C. Liquid SD-CAA cultures (1ml, composition: 20 g glucose, 6.7 g yeast nitrogen base, 5 g bacto casamino acids, 5.4 g Na_2_HPO_4_, 8.56 g NaH_2_PO_4_ per 1L) were inoculated by a single colony and grown overnight at 30°C (220 rpm). The grown cultures were spun down (3000 g, 3 min) and the culture media was replaced. The expression media - 1/9 media (18 g galactose, 2 g glucose, 8 g yeast nitrogen base, 8 g bacto casamino acids, 5.4 g Na_2_HPO_4_, 8.56 g NaH_2_PO_4_) was inoculated to OD 1.0 and cultivated at 30 °C overnight (12 – 14 hours, 220 rpm).

### Co-cultivation expression labeling and bait protein labeling procedures

According to the detection method, expression media was supplemented either by 1 nM DMSO solubilized bilirubin (Sigma Aldrich, USA) or purified 5 - 10 nM ALFA-tagged fluorescent protein (mNeonGreen) prior the culture cultivation. After the cultivation, cells were collected (3000 g, 3 min), washed once in ice-cold PBSB buffer, and subjected to analyses. Traditional antibodies based labeling procedure was done by using c-Myc Antibody (9E10, Cat # 626801, BioLegend, USA; incubation 1 h at 4°C) and Anti-Mouse IgG (Fc specific)-FITC antibody produced in goat (Cat # F4143, Sigma Aldrich, USA; 30 min at 4°C).

Bait proteins were labeled by amino-coupling fluorescence dye - CF® 640R succinimidyl ester (Biotinum, USA) according to the manufacturer’s protocol. Briefly, proteins were transferred to 100 mM bicarbonate buffer (pH 8.2) by using Amicon® Ultra Centrifugal Filters (3 kDa MWCO, Merck, USA) and mixed with 1: 3 ratio between protein and CF® 640R succinimidyl ester dye. The mixture was incubated in the dark at room temperature for 1h. After the incubation, the solution was transferred in GeBAflex-Midi Dialysis Tubes (8kDa MWCO, Geba, Israel) and dialyzed twice against 500 ml of PBS buffer at 4°C (8 – 12 h). The streptavidin conjugated with APC was purchased commercially (Cat#405207, BioLegend, USA).

### Cytometry analyses and FACS sorting

Expressed yeast cells were analyzed by using BD Accuri™ C6 Plus Flow Cytometer (BD Life Sciences, USA). The gating strategy is shown in Supplementary Figure S8. Green fluorescence channel (FL1-A) was used to record eUnaG2 or FITC signals representing expression positive cells, and a far-red fluorescent channel (FL4-A) recorded CF®640R stained proteins binding signals. No compensation was applied. Negative cells – EBY100 cells without plasmid or non-labeled cells were used to determine the negative population and set quadrant gating. Quadrants were used to divide the gated cell population into four plots showing negative (LL), non-specific (UL), expression (LR), and binding (UR) populations.

Fluorescence-activated cell sorting (FACS) experiments were done by using S3e Cell Sorter (BioRad, USA). Cells with surface-expressed proteins, detected *via* eUnaG2, DnbALFA co-cultivation labeling, or c-myc antibody labeling were incubated 1 h at 4 °C with bait protein and mixed by using lab rotator (5 rpm). Before the sorting, samples collected by centrifugation (3000 g, 3 min), 1 to 3 times washed with ice-cold PBSB buffer (1 ml) and passed through a cell strainer (40 μM, SPL Life Sciences, Korea).

### Binding assays and affinity curve determination using yeast display

Aliquots of expressed cells (10^6^) were collected (3000g, 3 min) and washed in PBSB buffer. The cell pellets were subsequently resuspended in analysis solutions across a range of concentrations. The composition of analysis solutions was as follows: PBSB buffer supplemented with a given concentration of ligand - CF®640R labeled bait protein (IL-28R1 or Kan-Nfr) and DMSO solubilized bilirubin (1 nM final concentration). The aliquots were incubated 1 h at 4 °C and mixed by using a lab rotator (5 rpm). Prior to the cytometry analysis, samples collected by centrifugation (3000 g, 3 min), 1 to 3 times washed with ice-cold PBSB buffer (1 ml) passed through a cell strainer (40 μM, SPL Life Sciences, Korea) and analyzed. The number of washes was increased depending on the background fluorescence. Usually, bait concentrations higher than 100 nM required multiple washing steps. Mean FL4-A values for expressing population subtracted by negative population FL4-A signals were used for determination of binding constant *K*_d_. The fitting of the standard non-cooperative Hill equation was done via nonlinear least-squares regression using Python 3.7. The total concentration of yeast exposed protein was fitted together with two additional parameters describing the given titration curve similarly to (Starr et al., 2020).

### Confocal fluorescence microscopy

Yeast cells were imaged with an Olympus FluoView FV1000 IX81 Spectral/SIM Scanner confocal laser-scanning microscope (Olympus GmbH, Hamburg, Germany), using 60 X phase-contrast oil-immersion objective, numerical aperture 1.35. The confocal sampling speed was set at 8 μs/pixel. The confocal aperture (C.A.) was fixed at 120 μm for all measurements. The images were collected at 640*640 (in pixel) or 52.4*52.4 (in μm). dimensions in line sequential mode. Yeast cell samples were put as liquid drops in glass slides with coverslips. To avoid evaporation of the sample, an extra cover glass was placed in the upside direction so that measurements of the samples could be done in a sandwich mode. Three channels were used for image collections: Fluorescence green channel (excitation at 488 nm and emission at 502–550 nm), fluorescence red channel (detecting the product formation, excitation at 559 nm and emission at 575–675 nm), and a third channel to visualize the transmission image. The laser at 488 nm was operated with 2-5 % of its maximum power and the laser at 559 nm was with 20% of its maximum power depending upon the sample expression qualities. For better image quality 4x to 8x zoom variations have been used.

## Acknowledgments

This research was funded by a grant from the Israel Science Foundation (ISF) number 1268/18 and by the United States – Israel Binational Science Foundation (BSF) number 2015376. We want to thank Ms Lucie Kolařová for providing us the IL-28R1 protein.

## Supplementary material section

### The step-by-step enhanced yeast display protocol

#### Cell maintenance, transformation, expression, and freezing procedures

1) Thaw a frozen aliquot of yeast cells and warm it rapidly up by hand. Remove the cryopreservant media by centrifugation (3000 g, 4 min), resuspend cells in an appropriate amount of YPD media, and let them recover overnight in a shaken incubator (30 °C, 220 r.p.m). The viable yeast will grow overnight to an absorbance of approximately 6 (OD600). This initial culture can be stored at 4°C for two weeks without the need for sub-culturing. Selective media (SD-Ura-his) can be used for *S.cerevisiae* EBY100 cells growth instead of rich media.
2) For the preparation of yeast electrocompetent or chemically competent cells follow the procedures described by Benatui et al (Benatuil et al., 2010) and Gietz et. al (Gietz and Schiestl, 2007; Gietz and Woods, 2002). Transformed cells bearing pJYD plasmids should be selected and maintained on tryptophan free media SDCAA/ SD-Trp. The viability and library size can be tested by serial dilutions on SDCAA/SD-Trp plates. Before expression cultures a liquid starter culture has to be prepared (overnight, 30 °C, 220 r.p.m).
3) Start the expression culture by pelleting 1 ml of starter culture cells (3000 g, 4 min). Remove the media and resuspend cells in 1/9 expression media to the OD 1. Cell expression conditions and temperatures may vary depending on protein and selection purposes (24 – 72 h, 20 – 37°C). Generally, proteins that are hard to express require lower starting OD (0.5 – 0.8), lower expression temperatures, and longer incubation times. The exact values have to be experimentally determined. Do not allow cells to reach the late stationary phase at which the surface expression drops, as protein quality may decrease due of media acidification. Cell co-cultivation labeling allowed in enhanced yeast display is described in detail in the chapter dedicated to labeling.
4) For *S. cerevisiae* EBY100 cells, electrocompetent cells, transformed cells and libraries freezing procedures follow the procedure described by Suga et al. (Suga et al., 2000) which we discover to be more efficient than traditional procedures based on glycerol or DMSO.

#### Co-cultivation labeling of yeast cells for enhanced yeast display

The labeling procedures of enhanced yeast display are versatile and depending on the plasmid used in the experiment. Here we will introduce the most common procedures utilizing the eUnaG2, DnbALFA and ALFA tag binding fusion proteins.

5) eUnaG2 reporter co-cultivation labeling procedures. Prepare the free-bilirubin (Sigma, SKU 14370) in DMSO solution (2 μM). This solution should be kept at −20 °C and can be used for a period of at least 6 months. Prolonged incubation at room temperature or in light will greatly reduce its stability. The solution is 2000 times concentrated stock and can be directly added at the inoculated expression culture. We do not recommend the preparation of 1/9 media with bilirubin since it compromises its stability. Expression culture without bilirubin can be used for different labeling purposes. The absence of bilirubin in cultivation media at the culture start can be replaced by its addition at least an hour before the harvesting or by 15 min labeling on ice with PBSB supplemented with 10 nM bilirubin. Bilirubin itself is not fluorescent in DMSO solution. A slight media color change is not compromising the fluorescence analysis.

The plasmid pJYDN3 is dedicated specifically for protein retention analysis: Two cultures should be expressed simultaneously – one with the addition of bilirubin (total eUnaG2 signal) and the second one bilirubin-free. The bilirubin-free culture surface expression is subsequently visualized by cell labeling procedure with PBSB supplemented with purified 5 mM eUnaG2-DnbALFA (cell-surface signal only, 30 min on ice, two washes with PBSB buffer). The same fluorescence signal reporter ensures easy extracellular/intracellular signal deconvolution without the need for fluorescence intensity calibration required with different fluorescent probes e.g. using antibody-based c-myc labeling.

6) DnbALFA reporter nanobody co-cultivation labeling procedures. The DnbALFA reporter enables any color labeling with either purified ALFA-tagged fluorescent proteins or fluorescently labeled peptides. The co-cultivation labeling has to be optimized for the labeling agent used. The optimal concentration for purified ALFA-tagged mNeonGreen was 5 nM. The co-cultivation labeling can be replaced by the addition of a labeling agent at least an hour before the harvesting or by 15 min labeling on ice with PBSB supplemented with 10 nM bilirubin.

#### Sorting process

7) The expression labeling strategy and the target labeling have to be compatible. We recommend using amino-reactive succinimidyl ester-based dye labeling instead of biotinylation and coupling with streptavidinAPC, especially for small target proteins to prevent false positive (streptavidinAPC binding) or false-negative results (sterical hindrance). Grow expression culture of yeast with the population size higher than 10-times the library complexity and wash cells once with PBSB buffer (3000 g, 4 min). Resuspend cells in an appropriate volume of PBSB buffer supplemented with the labeled bait protein. The volume and incubation time is dependent on the affinity (Chao et al., 2006) or selection strategy (Cohen-Khait and Schreiber, 2016). Larger volumes are generally needed to avoid bait protein depletion with concentrations lower than 1 nM. Incubate labeling mixture on rotator shaker to ensure homogenous binding conditions (5 rpm, 4 – 20°C). After bait protein incubation wash cells once with PBSB buffer (3000 g, 4 min). Keep cells pelleted prior to sorting to prevent dissociation.
8) Use a small sample of your cells to adjust your FACS sorter device setting and gating strategy for your experiment. If the negative (not expressing population) shows higher signals than expected, wash your cells again with PBSB buffer (3000 g, 4 min). Choose sorting gate in accordance with your selection approach – usually the double-positive quadrant. The general rule for sorting strategy is to start with a broad gating strategy (select 5 – 10 % of the population) and increase stringency in subsequent rounds up to the top 0.1 % of the double-positive population. This strategy prevents the loss of unique clones that did not express their properties due to competition with a large excess of other yeasts. We recommend sorting a minimum of 20k cells. After sorting, concentrate cells by centrifugation (3000 g, 4 min) and resuspend cells in an appropriate amount of SDCAA media and grow them for 24 – 48 h at 30°C (220 r.p.m). Repeat growth/sorting cycles until the population is substantially enriched.

#### Single-cell isolate characterization

9) After the population enrichment for the desired cells (three to five rounds of FACS) the verification of multiple single clones usually takes place before more in depth analysis. This step comprises of flow cytometry analysis that confirms binding to the selected target and sequencing of selected clones. The sequencing steps can be significantly simplified over the traditional methods which require enzymatic lysis, commercial yeast plasmid purification kits, *E.coli* transformation, and additional mini-prep purification (Chao et al., 2006).

We have discovered that the method for yeast chromosomal DNA isolation published by Lõoke et al. (Looke et al., 2011) can be modified for yeast plasmid preparations. Both plasmid preparations, which cover the whole enriched library and single clones can be prepared. The procedure consists of lithium acetate SDS cell lysis and ethanol precipitation. The resulting pellet contains total DNA (genomic and plasmid) from the sample and can be further subjected to a standard mini-prep isolation procedure. The obtained DNA is pure enough to be directly used in PCR applications, for transformations, or as a starting material for mini-prep procedures. In addition, the whole process of clone sequencing can be further accelerated by direct sequencing of colony PCR product, treated with ExoSAP enzyme mix (Bell, 2008).

Sequencing primers for pJYDN plasmids are: forward (GaL1b)
CCTCTATACTTTAACGTCAAGGAG, reverse (seq_R1)
CGGTGAAAATAGATGGGAACCTC.
Sequencing primers for pJYDC plasmids are: forward (C_seq_F)
GCAGCCCCATAAACACACAGTATG, reverse (pCT_seq_R)
CATGGGAAAACATGTTGTTTACGGAG.

#### DNA preparation note: Application of restriction-free cloning for targeted libraries

We took advantage of recent rapid developments in restriction-free cloning methods and applied them to improve and simplify our mutagenic workflow. We described in the main text, the construction of small libraries made of multiple *in silico* predicted mutations. The construction of these libraries was based on the work of Dr. Yoav Peleg’s lab (Erijman et al., 2011). Libraries were created by the multi-primer mutagenic PCR reaction. The principle of this method is based on the generation of different amplicons, so-called megaprimers, and their incorporation in the destination plasmid. A restriction-free approach coupled with error-prone PCR can be adopted in order to mutate selected region or regions within the gene of interest. The workflow consists of three subsequent PCR steps. In the first step, the random library is generated from the desired segment via error-prone PCR or similar methods. This amplified mutagenized fragment is used as a megaprimer in the next step. The second step is the restriction-free incorporation of library megaprimer in the destination vector. The reaction template is removed by *Dpn*I cleavage and purification. At last, the whole gene is amplified with recombination overhangs. This approach reduces the necessary steps compare to traditional methods like overlap-extension PCR and allow for semi-targeted mutagenesis which is difficult to achieve with different methods.

#### Media compositions

##### SDCAA (1 L)

20.0 g glucose
6.7 g yeast nitrogen base
5.0 g bacto-casamino acids
5.4 g Na_2_HPO_4_
8.56 g NaH_2_PO_4_

##### 1/9 expression media (1 L)

18.0 g galactose
2.0 g glucose
8.0 g yeast nitrogen base
8.0 g bacto-casamino acids
5.4 g Na_2_HPO_4_
8.56 g NaH_2_PO_4_

##### Amino acid composition of SD-W (1 L)

20 mg Adenine
20 mg Arginine
80 mg Aspartic acid 20 mg Histidine
30 mg Isoleucine
100 mg Leucine
30 mg Lysine
20 mg Methionine
50 Phenylalanine
200 mg Threonine
20 mg Tryptophane
30 mg Tyrosine
20 mg Uracil
150 mg Valine
6.7 g yeast nitrogen base

---

**Figure S1.**
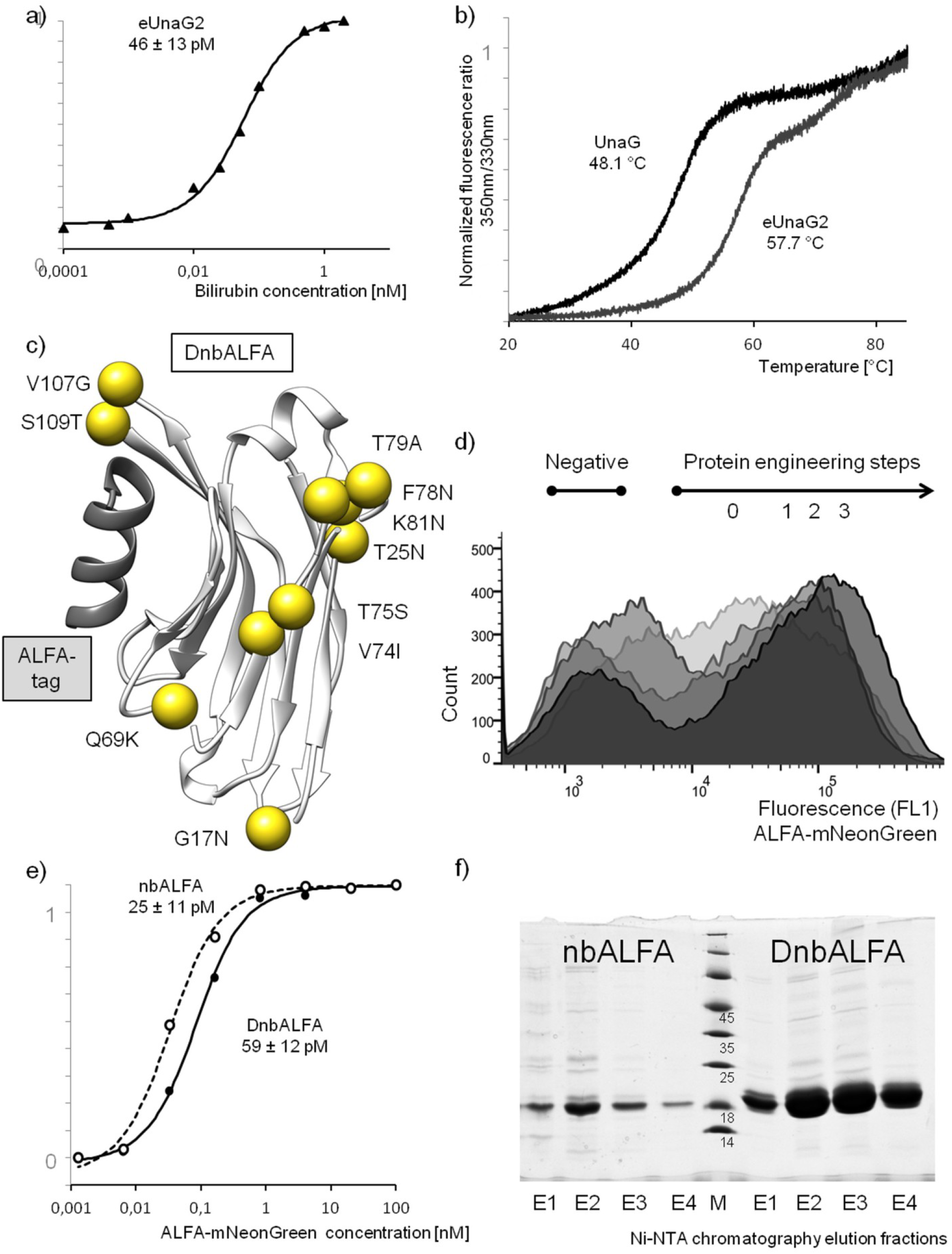
eUnaG2 properties and tailoring ALFA-tag binding nanobody for effective yeast display exposure by protein engineering. a) Bilirubin titration curve of eUnaG2. The affinity is higher than the affinity reported for UnaG (98 pM)(Kumagai et al., 2013) b) Melting of UnaG and eUnaG2 measured by using DSFnano Prometheus NT.48 instrument (Nanotemper Technologies). c) Mutations in designed DnbALFA introduced by protein engineering compared to original nbALFA (PDB id: 6I2G). d) Flow cytometry histograms showing the green fluorescence signal (FL1 channel) of different variants of nbALFA exposed on yeast surface and visualized by using Anti-c-myc antibody labeling. 0 – wild-type, 1 – Q69K, 2 – combining all predicted stabilization mutations, 3 – combining two N-glycosylation gaining mutations (G17N, T25N) with all stabilizing mutations; e) Binding curve between ALFA-tagged mNeonGreen and ALFA-tag binding nanobodies nbALFA and DnbALFA. f) SDS-PAGE analysis of expression of nbALFA and DnbALFA in 200 ml 2YT culture of *E.coli* BL21 (DE3) after single-step purification on Ni-NTA agarose. M – Protein ladder (kDa).

**Figure S2.**
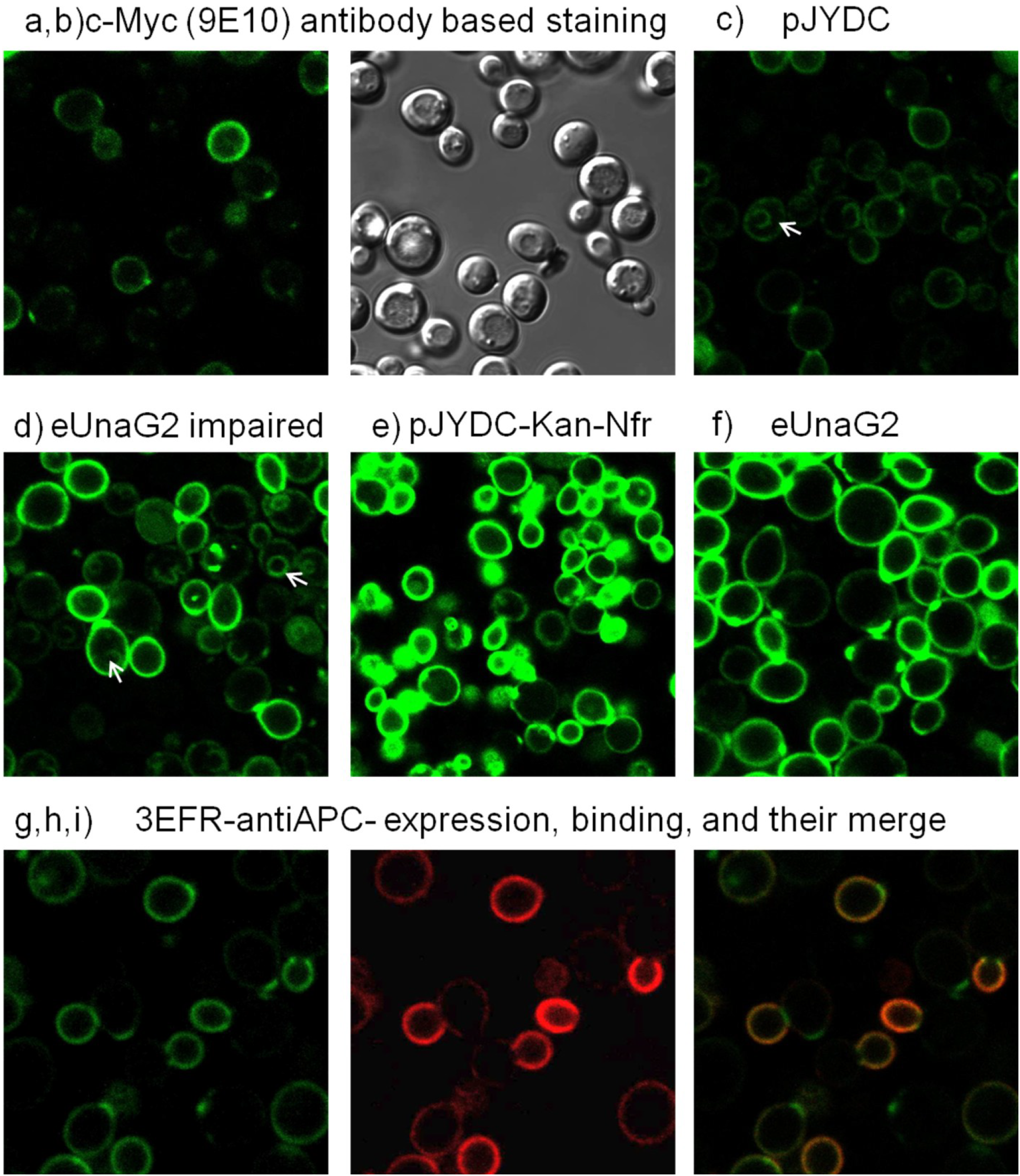
Microscopy analysis of protein expression and binding. a,b) The pJYDC1 expression analysis by using traditional c-myc labeling showed leaky expression of the pJYDC1 vector (media without bilirubin). c) pJYDC1 expression analysis using eUnaG2 fluorescence signal (media with 5 nM bilirubin, white arrow highlights the endomembrane system signal). The empty pJYDC1 vector-based expression resulted in the low fluorescent signal, which was predominantly located in the endoplasmic reticulum due to the presence of the HDEL endoplasmic reticulum retention signal sequence. d) pJYDC1-hUNG2 expression (UniProt: P13051). The hUNG protein was inserted in the pYJDC1 cleaved vector (*Nde*I and *Bam*HI) via homologous recombination. Although this process led to the expulsion of the HDEL sequence, the hUNG protein was not correctly processed through the secretory pathway and was partially retained probably in the ER (white arrows). e) pJYDC1 vector-based expression of Kan-Nfr wild type protein. The Kan-Nfr was inserted in the same position as hUNG but was correctly processed to the yeast surface. No signal from the endomembrane system was observed and the overall fluorescent intensity increased 4-times as measured by FACS. f) Positive control for eUnaG2 expression (pJYDNp plasmid). j,k,l) 3EFR-Cfr library (pJYDN) selected Streptavidin binding protein expression on yeast surface, StreptavidinAPC signal, and merged image.

**Figure S3.**
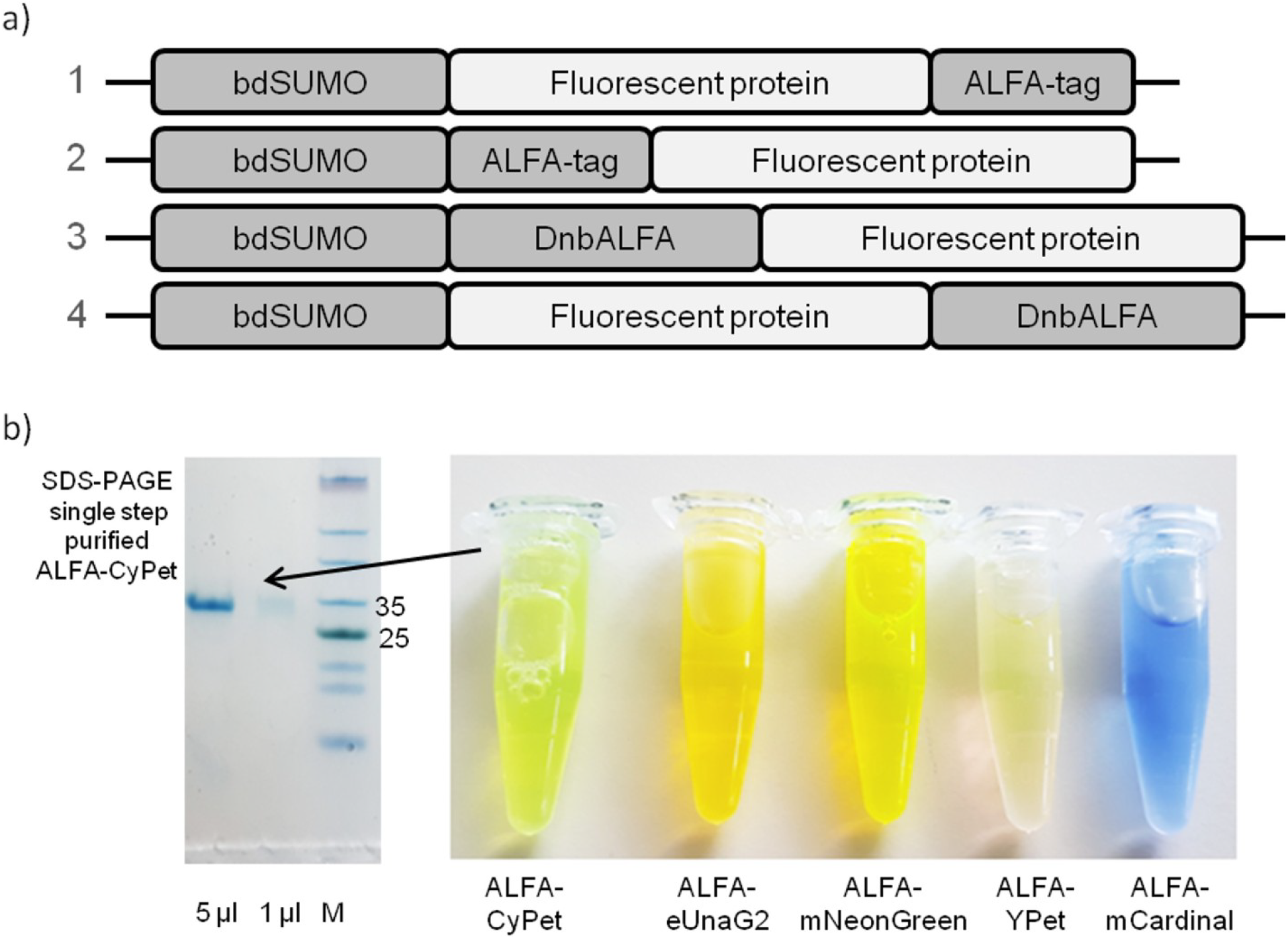
ALFA-tagged fluorescent proteins and fusions with DnbALFA for yeast display. a) Schematic representation of tested protein fusion constructs. Constructs 1 and 4 show a substantially higher yield of soluble fusion protein than constructs 2 and 3. b) Single-step purified proteins for use in pJYD yeast display plasmids. Proteins were expressed in 600 ml (2YT media) culture of *E.coli* BL21 (DE3) and purified by bdSUMO on-column cleavage.

**Figure S4.**
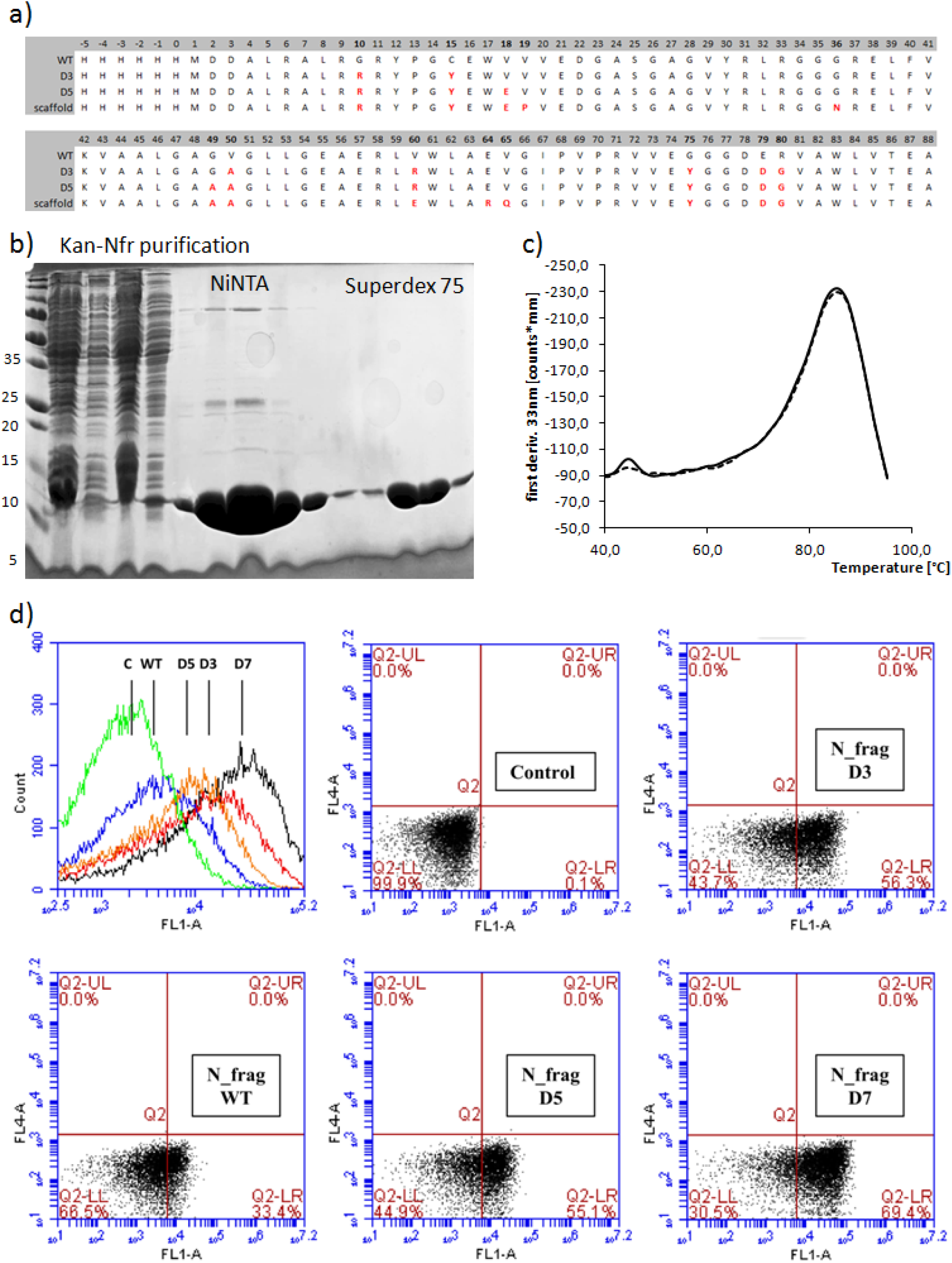
Sequence design and testing of Kan-Nfr scaffold. a) Sequence alignment among wild type, design stabilized 4H05 N-fragment 3 (D3), design stabilized 4H05 N-fragment 5 (D5) and final design stabilized 4H05 N-fragment 7 (D7 or Kan-Nfr scaffold). Mutated residues are highlighted in red. b) SDS-PAGE analysis of Kan-Nfr scaffold expression in *E.coli* BL21(DE3) and its two-step purification on NiNTA agarose and Superdex 75 16/600 gel filtration chromatography. c) Melting of Kan-Nfr measured by using DSFnano Prometheus NT.48 instrument (Nanotemper Technologies, duplicate). d) *S.cerevisiae* EBY100 cell surface expression of wild type and stabilized 4H05 N-fragments analyzed by BD Accuri™ C6 Plus Flow Cytometer.

**Figure S5.**
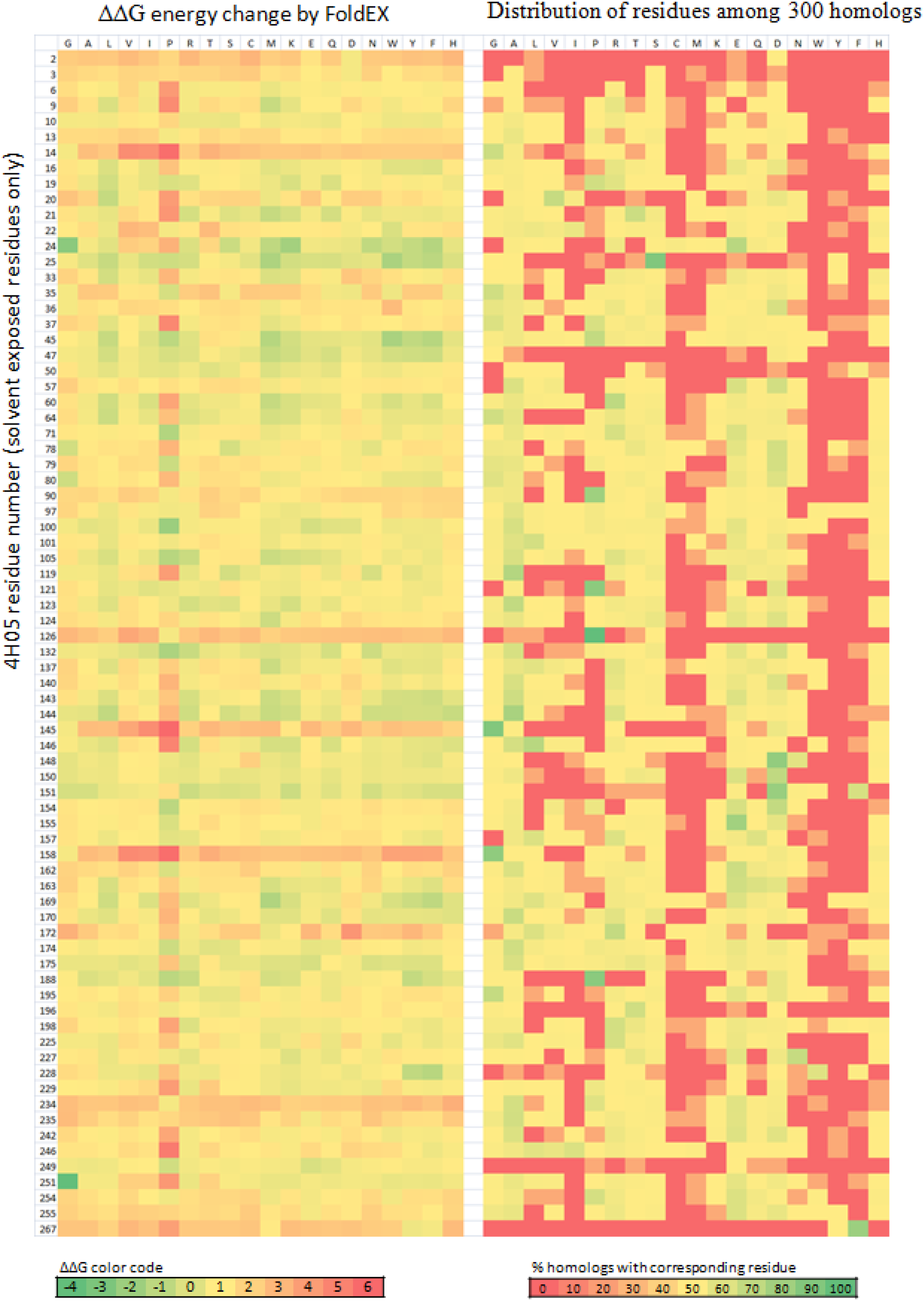
FoldEX and MSA comparison for mutation site identification in PDB: 4H05 (full protein). Only amino-acid residues with more than 50% solvent accessibility were evaluated. Positions and patches in protein structure with large evolutionary variability and narrow energy scale were searched. Multiple libraries were proposed covering the N-terminal region (residues −5 – 92) and the C-terminal part (93 – 267).

**Figure S6.**
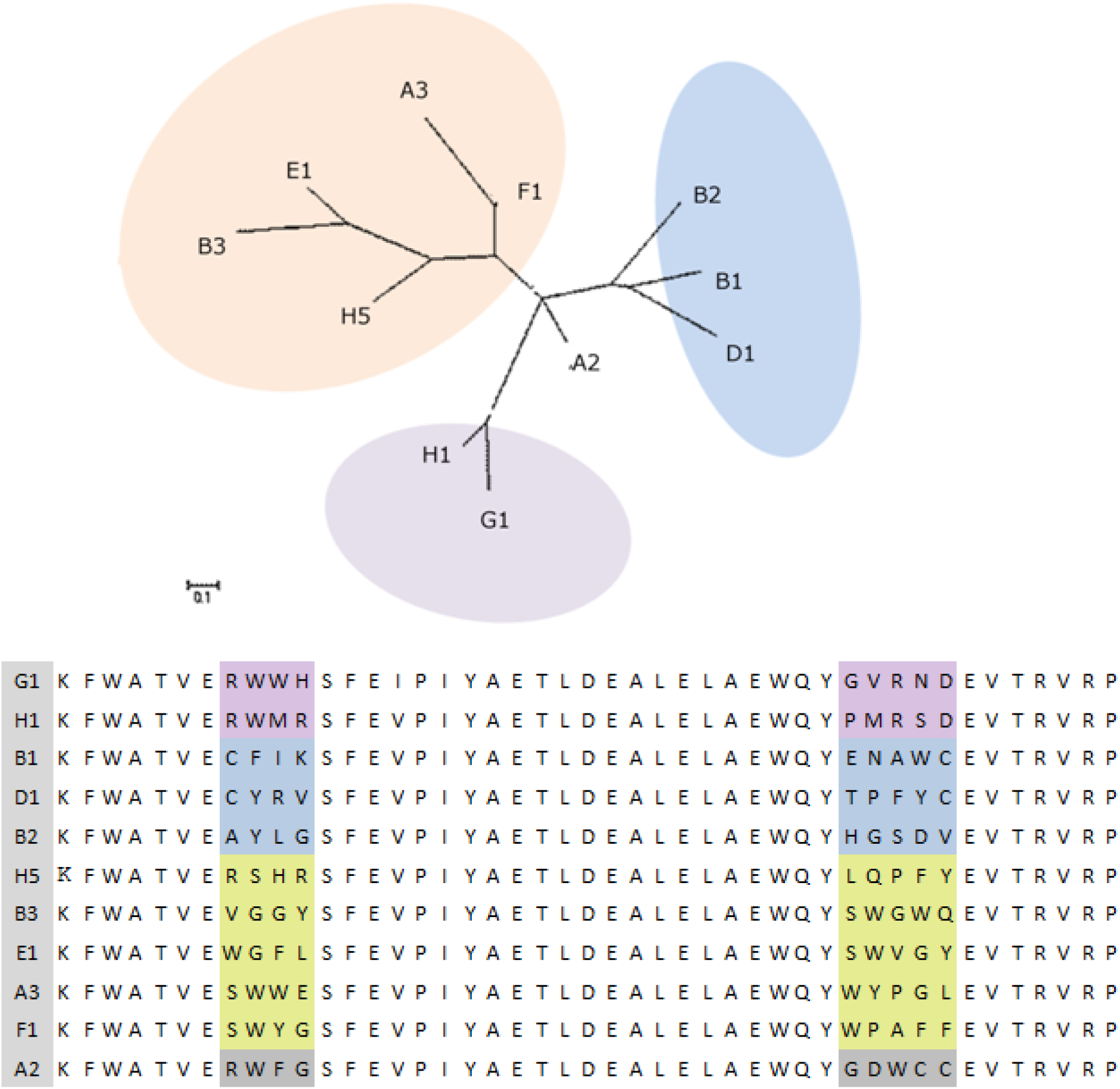
Neighbor-Joining Phylogeny of GP2 clones selected against Kan-Nfr protein. The phylogeny was constructed in Ugene software (http://ugene.net/) with JTT model.

**Figure S7.**
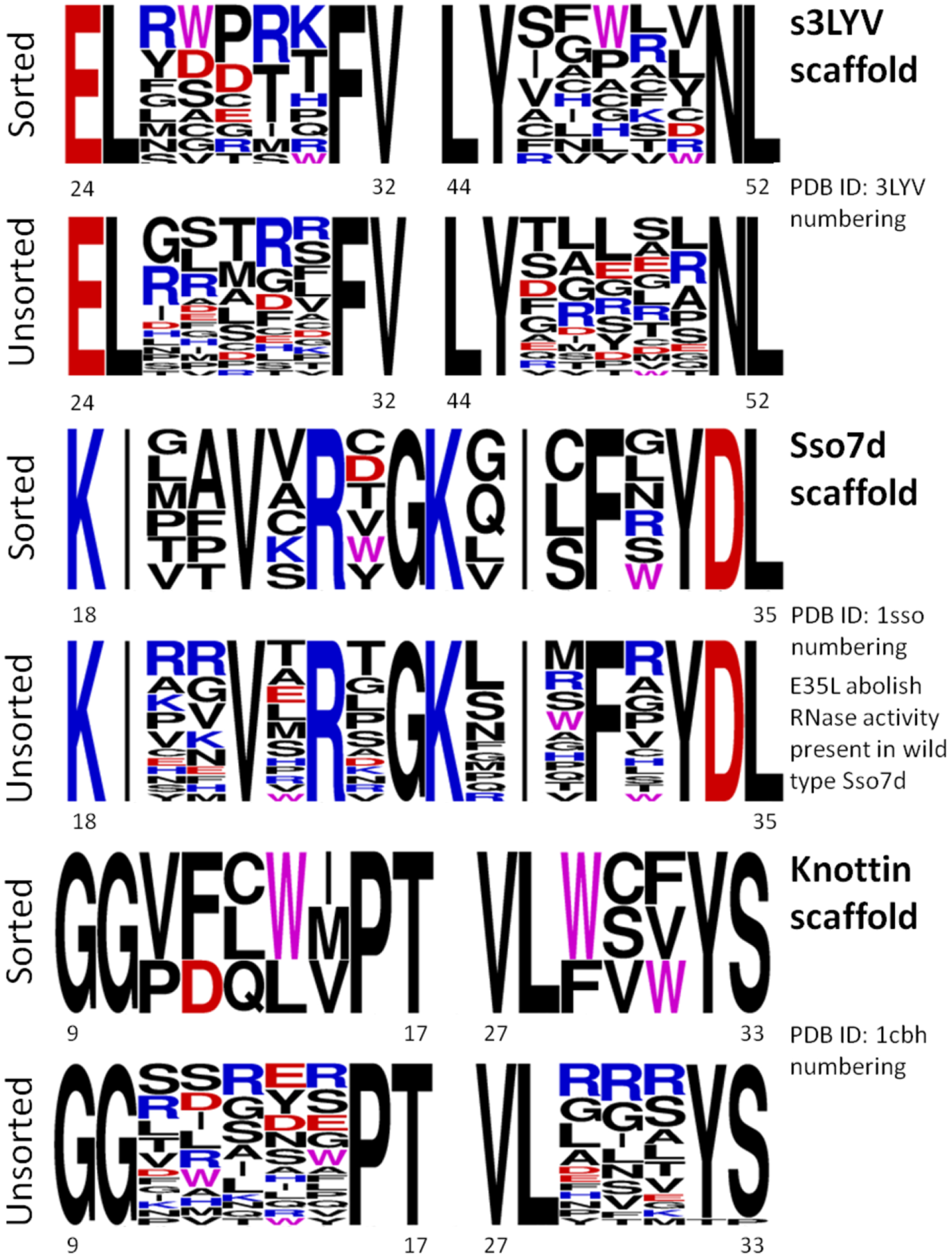
Outcome from low stringency selection among s3LYV, Knottin, and Sso7d libraries.

**Figure S8.**
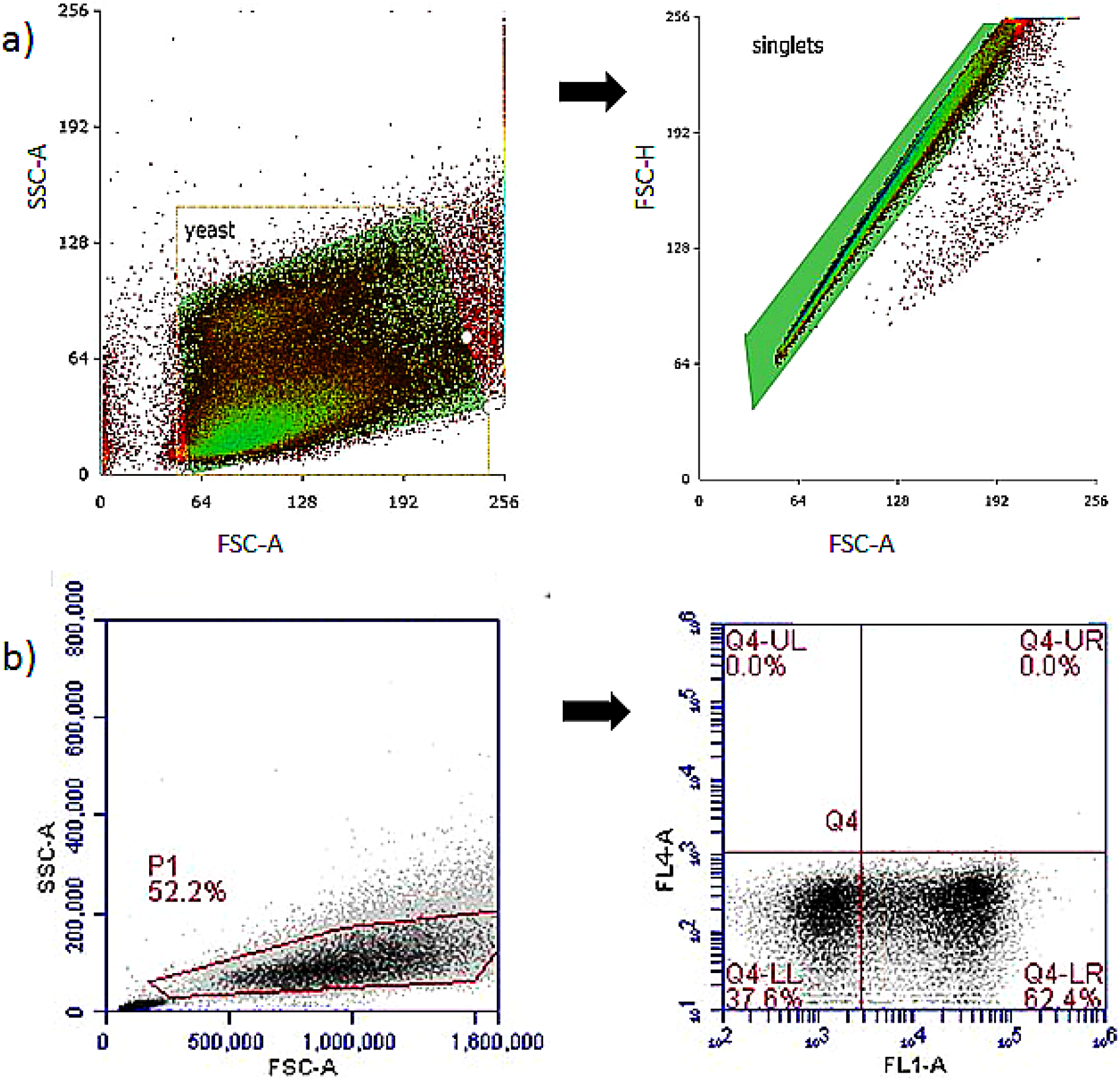
Gating strategy. a) Two-step gating strategy used in all sorting processes on S3e Cell Sorter device (Bio-Rad, USA). In the first step, yeast cells are isolated including mother and daughter populations. In the second gaiting, single cells are isolated by using FCS. b) Simplified single-step gating strategy used for analysis only on BD Accuri™ C6 Plus Flow Cytometer (BD Biosciences, USA). Quadrant gates were used to distinguish among negative cells (Q4-LL), cells positive for expression (eUnaG2 signal, Q4-LR), and binding/ double-positive cells (Q4-UR, not present in this example).

List of stabilizing mutations in Kan-Nfr compared to wild-type (PDB: 4H05 structure numbering): G10R (randomized position), C15Y, V18E, V19P, G36N, G49A, V50A, V60E, E64R, V65Q, G75Y, E79D, R80G;

The comparison of 4H05 wild-type and scaffold sequence is shown in Figure S10a

List of stabilizing mutations in 3EFR-Cfr compared to wild-type (PDB: 3efr structure numbering): E190R, L194I, K198E, K202I, K209N (randomized position), L215E, E217D;

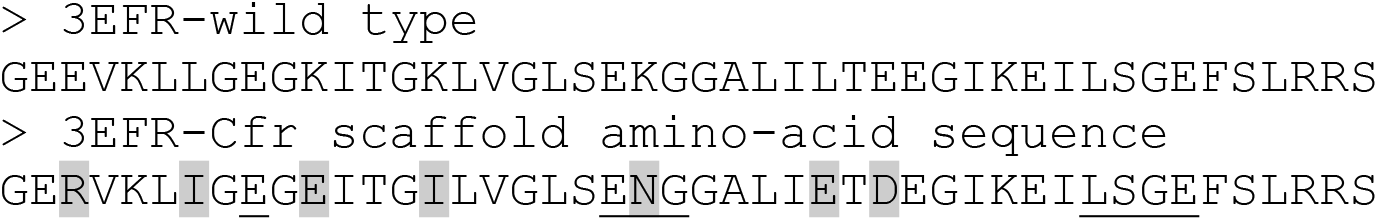

List of stabilizing mutations in s3LYV compared to wild-type (PDB: 3lyv:A structure numbering): V16S, I32V, D38T, A40Q, T41V;

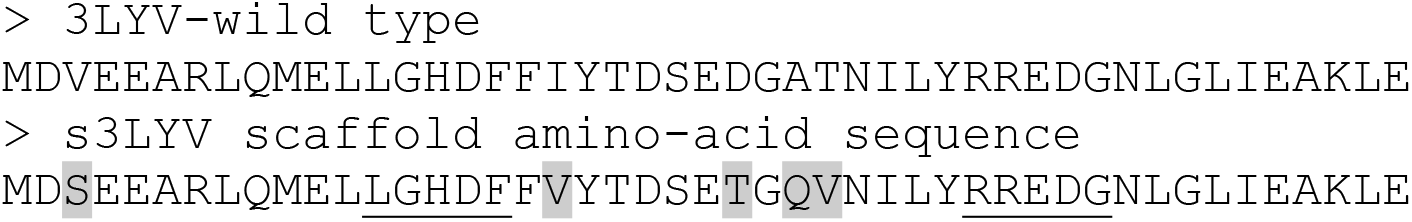

**Material S1 – List of stabilizing mutations incorporated in protein scaffold sequences**. The highlighted positions in amino-acid sequences correspond to mutant residues. The randomized residues are underlined.

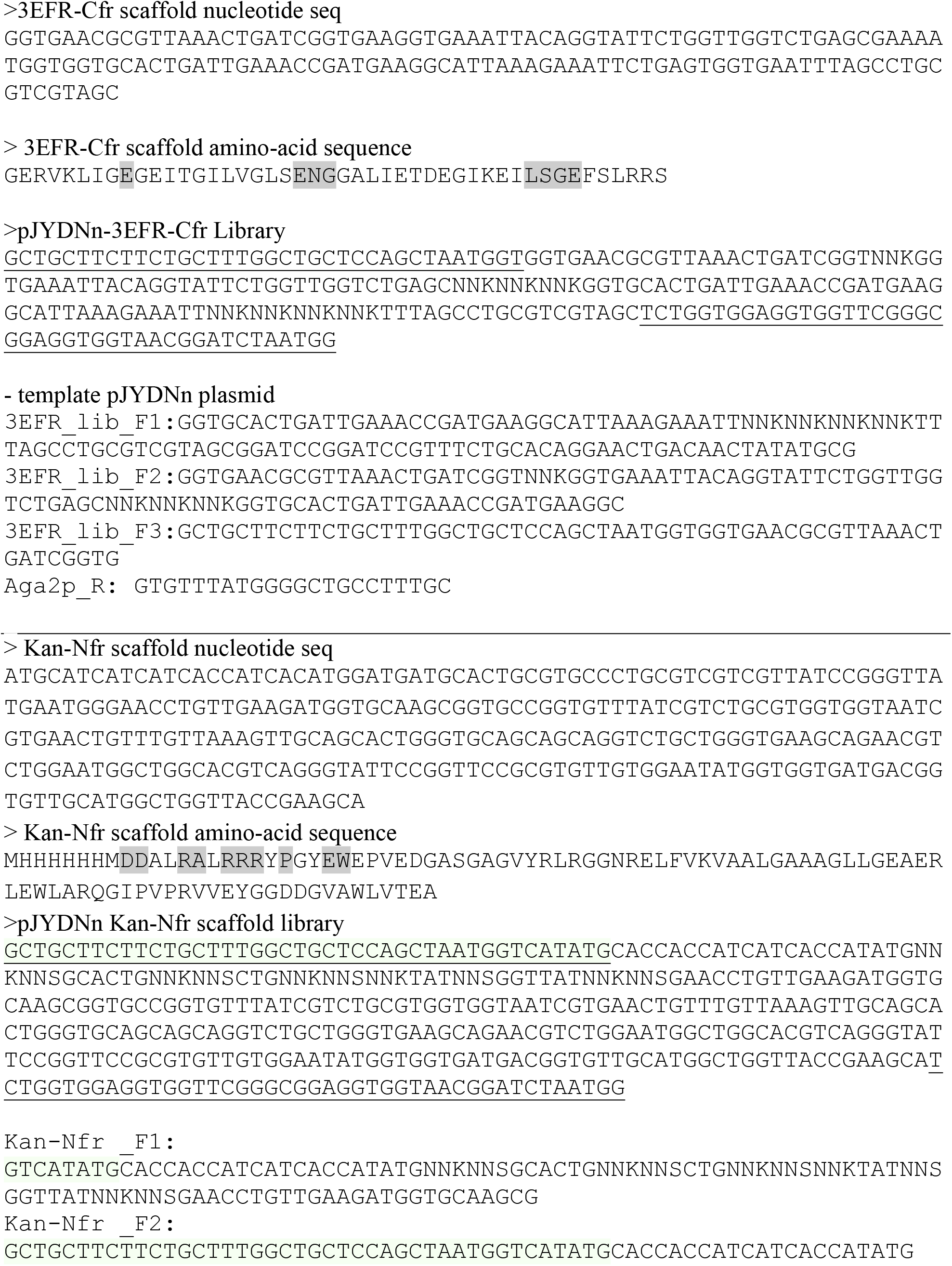
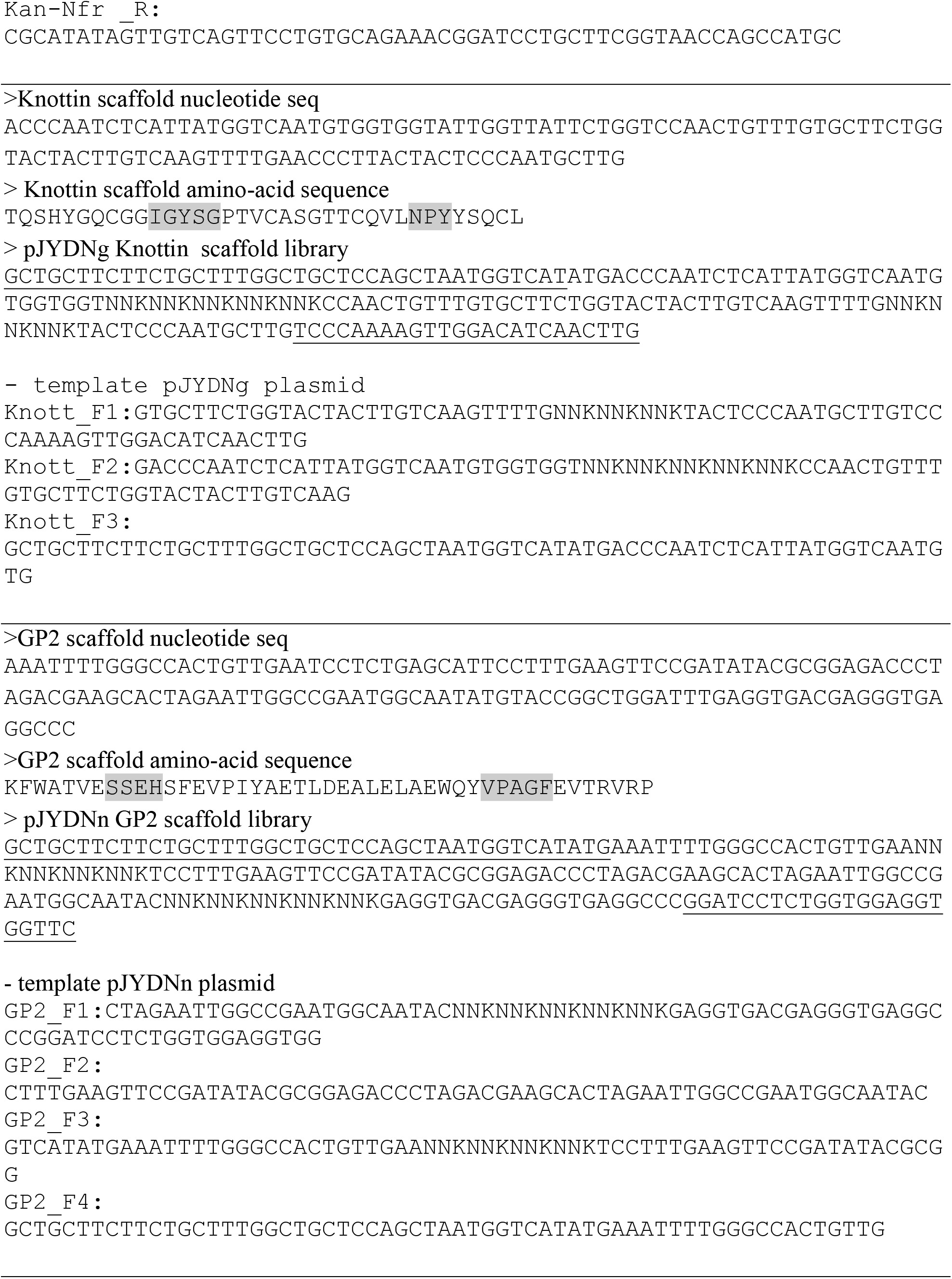
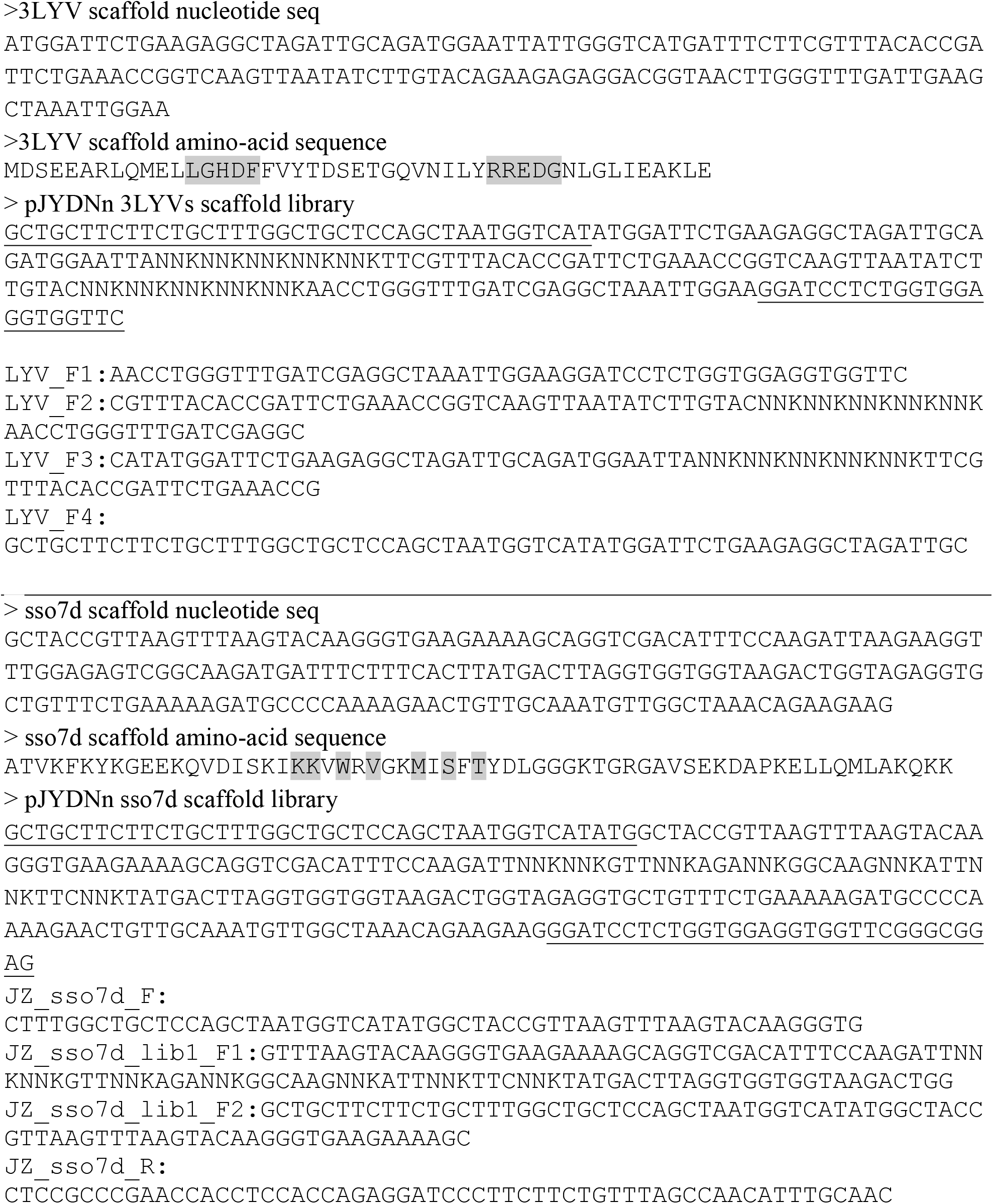

**Material S2 – Scaffold sequences, randomized positions, and final library constructs.**All scaffolds were optimized for expression in *S.cerevisiae*. The first nucleotide sequences were used for scaffold expression verification and also for control expression in *E.coli*. The amino-acids highlighted by grey background were randomized. Libraries sequences are including overhangs for homologous recombination (underlined). Some of them differ in non-randomized nucleotides compared to nucleotide sequences only. This difference is only on the codon usage level and was incorporated during library construction via PCR reactions.

